# Epigenomic anomalies in induced pluripotent stem cells from Alzheimer’s disease cases

**DOI:** 10.1101/2024.08.29.610372

**Authors:** Anthony Flamier, Alisar Katbe, Dounya Serhani, Rimi Hamam, Ryan Hogan, Erika Tavares, Élise Héon, Roy Hanna, Gilbert Bernier

**Affiliations:** Stem Cell and Developmental Biology Laboratory, Hôpital Maisonneuve-Rosemont, 5690 Boul. Rosemont, Montréal, Canada, H1T 2H2; Centre de recherche Azrieli du CHU Sainte-Justine, Montréal, Canada; Genetics and Genome Biology, The Hospital for Sick Children, Toronto, Canada; Department of Ophthalmology and Vision Sciences, The Hospital for Sick Children, Toronto, Canada; Institute of Medical Science, The University of Toronto, Toronto, Canada; Department of Neurosciences, Université de Montréal, Montréal, Canada; McGill University, Montréal, Canada

**Author notes:** Senior author (Roy Hanna).

**Keywords:** BMI1, Polycomb, Alzheimer, iPSC, RNA-seq, ChIP-seq Running title: Epigenome anomalies in Alzheimer iPSCs

## Abstract

Reprogramming of adult somatic cells into induced pluripotent stem cells (iPSCs) resets the aging clock. However, primed iPSCs can retain cell-of-origin epigenomic marks, especially those linked to heterochromatin and lamina-associated regions. Here we show that iPSCs produced from dermal fibroblasts of late-onset sporadic Alzheimer’s disease (AD) cases retain epigenomic anomalies that supersede developmental defects and neurodegeneration. When compared to iPSCs from elderly controls, AD iPSCs show reduced *BMI1* expression, lower H3K9me3 levels, and an altered DNA methylome. Gene Ontology analysis of differentially methylated DNA regions (DMRs) reveals terms linked to cell-cell adhesion and synapse, with the cognitive resilience-associated MEF2 family of transcription factors being the most enriched at DMRs. Upon noggin exposure, AD iPSCs show lesser efficient neural induction and forebrain specification, together with increased ZIC2, ZIC5 and WNT-related gene expression. Long-term AD neuronal cultures present a dedifferentiation and loss-of-cell identity phenotype. Despite these epigenomic anomalies, AD iPSCs generate cortical neurons in normal proportion and readily form cerebral organoids developing amyloid and Tau pathology. BMI1 overexpression in AD neurons mitigates amyloid and tau accumulation, heterochromatin fragmentation, and G4 DNA induction. These findings implicate reprogramming resistant epigenomic anomalies or uncharacterized genetic alterations working in trans on the epigenome in AD pathophysiology.

## INTRODUCTION

Late-onset sporadic Alzheimer’s disease (AD) is the most common dementia with a predicted prevalence in 2050 of 13.8 million in the United States and 152 million worldwide ^1,2^. The brain changes in AD begin 20 or more years before symptoms appear ^3^. 11% of people aged 65 and older have AD, and this prevalence grows to 32% for people aged 85 and more ^4^. At the genetic level, carriers of the *Apolipoprotein E (ApoE) epsilon 4* allele have a greater risk to develop AD ^5^. AD is characterized by progressive memory and behavioral impairment owing to degeneration of limbic and cortical areas of the brain. Pathological hallmarks of the disease are extra-cellular amyloid plaques and intra-neuronal phospho-Tau tangles formation, synaptic atrophy and neuronal cell death^6^. Amongst patients with amyloid plaques and Tau-tangles, a small fraction of those can maintain normal cognition, a phenomenon referred to as cognitive resilience ^7,8^. Recently, the Myocyte Enhancer Factor 2A and C (MEF2A/C) genes were identified as key players at promoting cognitive resilience in AD through activation of synaptic transmission and cognition-related genes ^9^. Chromatin profiling studies of AD brains also revealed an Alzheimer’s resilient epigenetic signature with a Gene Ontology (GO) for cell-cell adhesion via plasma membrane adhesion molecules, synaptic transmission, and neural development in neurons, and inflammatory response in microglia^10^. While the respective contribution of genetic, lifestyle, and environment in AD’s etiology remains complex, advanced age represents the greatest risk factor to develop AD ^4^. Aging is characterized by the accumulation of genetic mutations and loss of epigenetic information ^11,12^. In mice, faithful repair of genetically induced DNA breaks is sufficient to trigger accelerated aging, including erosion of the epigenetic landscape and loss of cell identity ^13^. Notably, single-cell analyses revealed epigenetic erosion and loss of cell identity in late-stage AD brains, a phenomena not seen in old control brains ^14^. AD brain’s neurons also show features reminiscent of accelerated aging, including loss of heterochromatin, alterations of the nuclear envelope, and persistent DNA damage response ^15–18^.

Proteins of the Polycomb Group (PcG) family form large molecular complexes that regulate epigenomic cell memory and play multiple functions during development, cancer, and aging ^19^. EZH2 is the catalytic unit of the Polycomb repressive complex 2 (PRC2) and mediates gene silencing through bi- and tri-methylation of histone H3 at lysine 27 (H3K27^me3^) ^20,21^. RING1 is the catalytic unit of the Polycomb repressive complex 1 (PRC1) and mediates gene silencing through mono-ubiquitylation of histone H2A at lysine 119 (H2A^ub^) ^22^. During development, one of the general functions of PcG is to prevent the expression of lineage-restricted genes in pluripotent stem cells, and later to prevent expression of pluripotency genes in somatic cells ^23,24^. The B-cell Moloney leukemia insertion site 1 (BMI1) protein prevents RING1a/b auto-ubiquitylation and degradation and works as the core unit for canonical PRC1 activities in somatic cells ^25^. The repressive histone mark H3K9^me3^ accumulates at gene-poor and repeat-containing heterochromatin ^26^. Notably, BMI1 is enriched at H3K9^me3^-associated heterochromatin and its deficiency impairs histone H3K9^me3^ deposition and maintenance ^27,28^. BMI1 is required to prevent senescence or apoptosis of somatic cells and cancer cells ^29–31,19^. *Bmi1*-null mice present growth retardation and cerebellar degeneration, together with premature ocular and brain aging ^32,33^. *BMI1* expression is reduced in AD brains, and *BMI1* inactivation in neurons results in the accumulation of beta-amyloid and Tau tangles ^34^. *Bmi1^+/-^* mice develop with old age behavioral and brain anomalies resembling AD ^15^. BMI1 protein levels are reduced in the serum of minimal cognitive impairment and AD patients ^35^, and a minor allele of *BMI1* was associated with slower cognitive decline in AD ^36^, altogether suggesting that BMI1 is an AD modifying gene.

The induced pluripotent stem cell (iPSC) technology allows reprogramming of somatic cells into pluripotent stem cells capable of self-renewal and differentiation into cell progenies of the 3 primordial germ layers ^37,38^. By analogy with sexual reproduction, reprograming rejuvenates adult somatic cells, erasing aging-associated epigenomic marks ^39^. Primed human iPSCs are developmentally equivalent to the post-implantation epiblast ^40,41^. While iPSCs and embryonic stem cells are apparently very similar, their epigenome differs substantially. Notably, iPSCs can acquire aberrant DNA methylation (DNAme) states and retain part of the epigenetic memory of the cell of origin, especially those at H3K9^me3^-enriched and lamina-associated chromatin regions ^42,43^. Similarly, mouse embryos produced by nuclear transfer retain from donor cells reprograming resistant chromatin regions enriched for H3K9^me3^ and which imped normal development ^44^. Multiple iPSC lines have been generated from AD patient’s somatic cells. Notably, differentiation of AD iPSCs into cortical neurons can recapitulate AD-related pathologies, such as elevated Aβ42 secretion, p-Tau accumulation, dendritic atrophy, and neuronal cell death ^34,45,46^. Interestingly, induced neurons produced by transcription factor overexpression in AD human dermal fibroblasts (HDFs) show reduced neuronal maturation, activation of stress response genes, and increased expression of stem cell and cancer-related genes. This phenotype was not observed in induced neurons produced from HDFs of old control donors ^47^. Since most AD patients do not carry defined disease-causing genetic mutations ^48^, it remains unclear however why AD neurons generated from either HDFs or iPSCs present these anomalies.

Herein we describe the molecular characterization of control and AD iPSC lines. We found subtle but significant gene expression differences between undifferentiated control and AD iPSCs, together with heterochromatin and DNAme alterations. Unexpectedly, differentially methylated DNA regions (DMRs) in iPSCs were highly enriched for MEF2s binding sites. Differentiation of AD iPSCs into neuroepithelial (NE) cells resulted in less efficient neural induction and elevated ZIC2, ZIC5 and WNT-related gene expression. Advanced AD neuronal cultures presented a dedifferentiation phenotype, expressed non-neuronal lineage genes, and developed several pathologies observed in AD brains. Many of these neuronal pathologies were improved following BMI1 overexpression.

## RESULTS

### Altered transcriptome and histone modifications in undifferentiated AD iPSCs

HDFs from elderly non-demented controls and AD cases were reprogram into iPSCs using plasmid vectors expressing the Yamanaka factors. Polymerase Chain Reaction analysis on genomic DNA confirmed the absence of plasmid vector integration after 10 passages (**Figs. 1A-C and S1A-D**) ^38,49^. The iPSCs showed comparable expression levels with the H9 human embryonic stem cell line for *POU5F1* (encoding for OCT3/4) and *NANOG*, but lower levels for *ZFP42* (REX01) (**Fig. S1E**) ^50^. CTL1, CTL2, AD2, and AD2 iPSC lines presented a normal karyotype and were able to generate teratomas in NOD/SCID mice (**Fig S1F-G**) ^38^. Whole genome sequencing (WGS) revealed that CTL1, CTL2, CTL14, CTL16, AD1 and AD2 iPSC lines were *APOE e3/e3* or *e2/e3* carriers and did not present mutations in protein-coding genes, including pathogenic variants in *APP*, *PSEN1*, *PSEN2*, *SORL1*, or *MAPT* (**Fig. 1C, and Supplementary Materials and Methods**). To account for variabilities in reprogramming methods, manipulation techniques, and cell culture conditions, we included a previously characterized AD iPSC line (AD3) generated using Sendai’s virus ^45^ and an independently reprogrammed cell line (AD17) from the same AD case as the AD1 cell line (**Fig. 1C**). We performed RNA-sequencing (RNA-seq) on the CTL and AD iPSC lines. Principal component analysis (PCA) revealed that CTL and AD iPSCs were similar in their heterogenicity (**Fig. 1D**), with no differences in the expression of pluripotency genes (**Fig. 1E**). Unsupervised cluster analysis revealed low but significant differences between CTL and AD iPSCs for ∼200 transcripts, many corresponding to non-coding RNAs and pseudogenes, with the top 100 most differentially expressed genes (DEGs) presented in **Fig. 1F**. From these, lower expression was found in AD iPSCs for *BMI1* and *KDM6A*. *KDM6A* encodes a H3K27-demethylase that antagonizes PRC2 activity ^51^. Notable DEGs up-regulated in ADs were *DYRK3*, *FOXK2*, *DNAJB1* (*Hsp40*), *HSPA1B* (*Hsp70-1B*), *HSPA1A* (*Hsp70-1A*), *NFKB inhibitor delta* (*NFKBID*), *JUNB*, and *MT1F* (**Fig. 1F**). DYRK3 associates with Hsp90 to promote stress granule dissociation and reinstate mTOR signaling ^52^. *FOXK2* control the unfolded protein response ^53^. Hsp40 and Hsp70-1A/1B are chaperones preventing protein misfolding under stress condition. The NFKB pathway plays a pivotal role in microglia and during neuroinflammation in AD ^54^. JUNB is involved in immune response and cancer ^55^. *MT1f* encodes a Zn and Cu metal ion and ROS response protein ^56,57^. DEGs in AD iPSCs are thus generally involved in immune signaling and cellular stress response.

**Figure 1.**
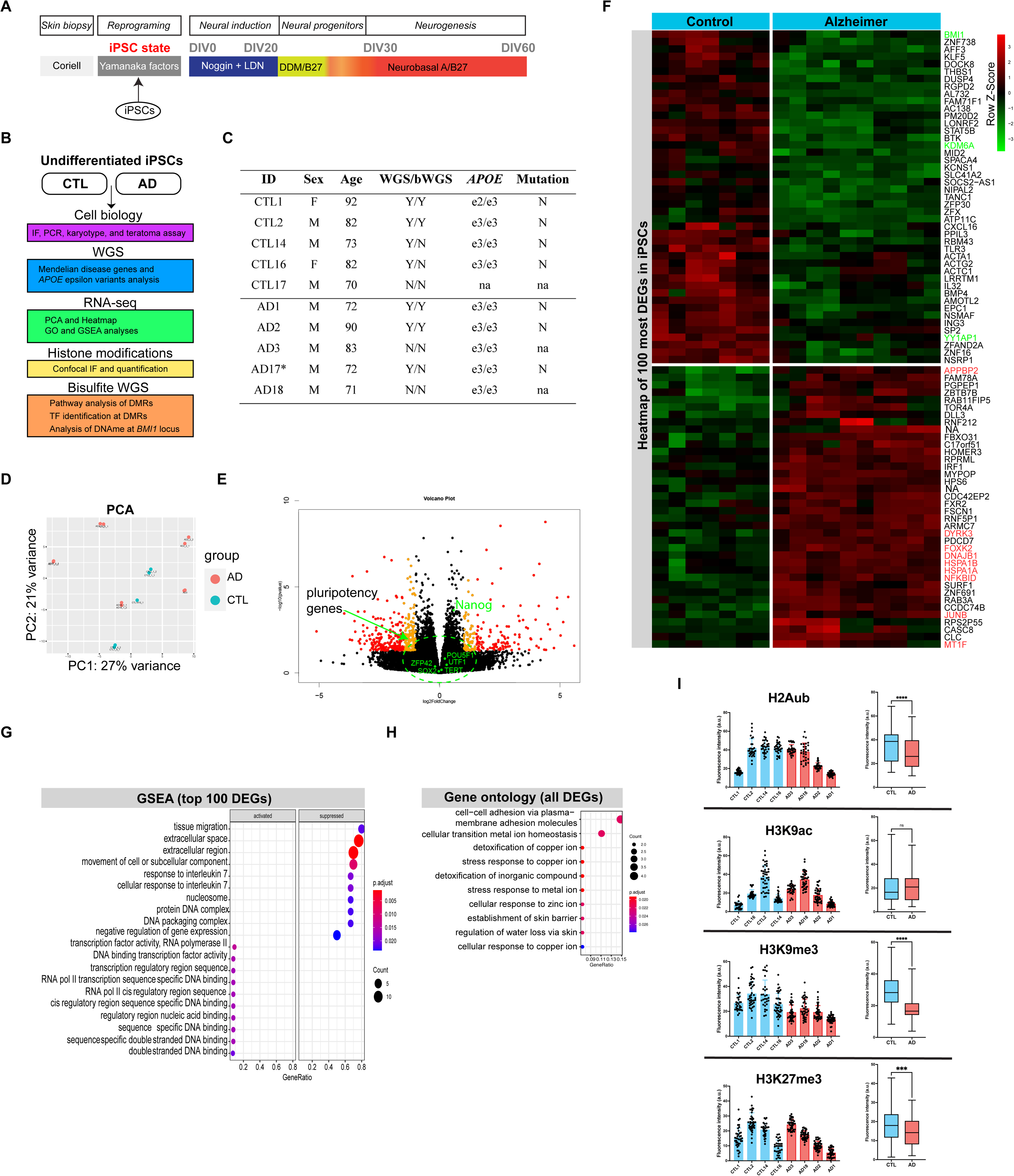
Altered transcriptome and histone modifications in AD iPSCs. (A) Schematic of the cell’s differentiation steps used in this study (iPSC stage). (B) Graphical representation of analyses performed on iPSCs. (C) List of iPSC lines, Whole genome sequencing (WGS), bisulfite Whole genome sequencing (bWGS), Yes (Y), No (N), not available (na). (D) Principal component analysis (PCA) of iPSC lines, with CTL = 4 cell lines with 7 replicates; AD = 4 cell lines with 10 replicates. (E) Volcano plot of all genes expressed in iPSCs (CTLsvs AD). CTL with n = 7 and AD with n = 10). Pluripotency genes are labelled in green. Significant DEGs at 2 folds (yellow dots) and more than 2 folds (red dots). (F) Heatmap of unsupervised clustering of top 100 DEGs between CTL and AD iPSC lines, with a cutoff of at least 2 folds. (G) GSEA of top 100 DEGs between CTL and AD iPSC lines. (H) Gene ontology analysis of top 100 DEGs between CTL and AD iPSC lines. (I) Quantification of histone modifications between CTL and AD iPSC lines (n = 30 nuclei across 3 independent pictures per cell line). All values are mean ± SEM. (***) P < 0.001 and (****) P < 0.0001 by Welch’s unpaired t-test.

Gene set enrichment analysis (GSEA) showed that tissue migration, extracellular space, movement of cell or subcellular component, response to interleukin 7, and nucleosome/DNA packaging complex/negative regulation of gene expression were suppressed in ADs (**Fig. 1G**). On the other hand, transcription factor activity related to RNAPol2 was slightly activated in ADs (**Fig. 1G**), suggesting the possibility of a more relaxed chromatin. Gene ontology (GO) analysis of DEGs revealed that cell-cell adhesion via plasma-membrane adhesion molecules (ranked #1), cellular transition metal ion homeostasis, and stress response to copper ion were the most prominent pathways (**Fig. 1H**). To investigate chromatin state, we performed immunofluorescence (IF) analysis of iPSC colonies using antibodies against the H2A^ub^, H3K27^me3^, H3K9^me3^ and H3K9^ac^ histone marks. We observed variations for all 4 markers between iPSC lines within the same group and between the CTL and AD groups. Quantitative analysis showed that all 3 repressive histone marks were reduced in ADs, while H3K9^ac^ levels were unaffected (**Fig. 1I**). However, most consistent results were observed for H3K9^me3^ because showing the lowest variance within groups (**Fig. 1I**). We concluded that AD iPSCs present subtle, but significant gene expression variations linked to cell-cell adhesion and stress response pathways together with alterations in histone modification suggestive of a more relax heterochromatin.

### Altered DNA methylome in AD iPSCs

To investigate for other epigenome alterations, we performed bisulfite WGS in CTL1, CTL2, AD1 and AD2 iPSC lines to identify differentially methylated DNA regions (DMRs). We found that AD iPSCs presented generally more DNAme at CHH and CHG dinucleotides when compared to CTLs before, after, and within gene’s body (**Fig. 2A**). While overall DNAme levels at CG was comparable between CTLs and ADs (**Fig. 2A**), AD iPSCs presented less hypermethylated CG DRMs (**Fig. 2B**). DNAme levels were also found comparable at LINE and LTR repeats (**Fig. S2A**). We annotated all genes showing DMRs between CTLs and ADs. Of these, 3 genes showed common DMRs for GC, CHH and CHG dinucleotides and were associated with human diseases i.e. SYTL1, HDAC11, and GSTM5 (**Fig. 2C**). GO analysis of all DMR-associated genes revealed that most were related to neuronal development, axonogenesis and synaptic transmission (**Fig. 2D**). Cell-cell adhesion and homophilic cell adhesion via plasma membrane associated molecules (ranked #1) was the most significant altered biological pathway for hypo- and hyper-DMRs with CG dinucleotides (**Fig. 2D**), similarly as observed in the RNA-seq dataset (**see Fig. 1I**). These GO terms are also near identical those associated with the Alzheimer’s resilient epigenetic signature ^10^. We curated all DMRs for transcription factor (TF) binding sites using JASPAR (10.1093/nar/gkab1113https://pubmed.ncbi.nlm.nih.gov/34850907/), revealing 235 commonly share TFs for DMRs bearing CG and CHH dinucleotides, but almost none for DMRs bearing CHG dinucleotides (**Fig. 2E**). Most commonly found TFs and TF families were ZNF460, ZNF135, MEF2A-D, NR2F1, PITX2 and SP1 (**Fig. 2E and F**). ZNF460 and ZNF135 encode zinc finger proteins with unknown function. The MEF2 family encode transcription factors involved in development, synaptogenesis and higher cognitive function, and showing neuroprotective activity in AD mouse models ^9,58,59^. Pathogenic *MEF2C* and *NR2F1* variants are associated with severe neurodevelopmental and autism spectrum-like disorders ^60–62^ ^63^. *PITX2* variants are linked to stroke and higher AD risk ^64^. *SP1* has been associated with AD and PSP-associated genes in hub-based models ^65,66 67^. We also analyzed the *COMMD3-BMI1* locus for the presence of DMRs. No DMRs were found between CTL and AD iPSCs (**Fig. S2B**). Methylated CG within CpG islands can be maintained through cell division and negatively influence gene expression ^68^. We found that two CpG islands within and downstream of the *COMMD3-BMI1* locus showed higher DNAme for CHH and (#53) and for CG and CHH (#225) in ADs (**Fig. 2SB**). Another CpG island (#134) located far upstream of the *COMMD3-BMI1* locus also showed higher DNAme for CG dinucleotides in ADs (**Fig. S2B**). Although correlative, this observation could explain reduced *BMI1* expression in AD iPSCs. We concluded that AD iPSCs thus present an altered DNA methylome with GO terms associated with cell-cell adhesion and synapse and showing strong enrichment for MEF2s at DMRs.

**Figure 2.**
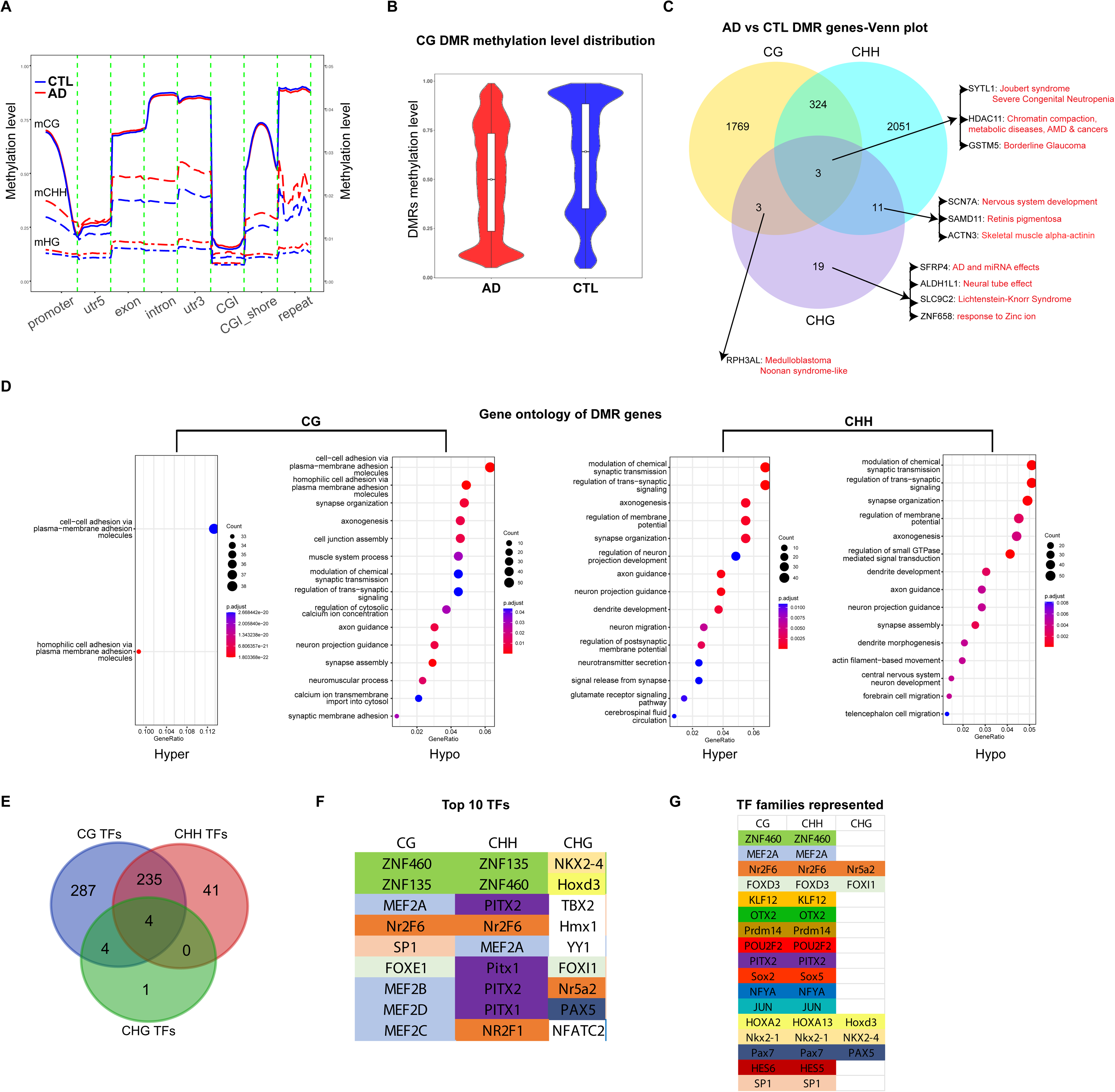
Altered DNAme in AD iPSCs. (A) Diagram showing global DNA methylation level between CTL1/CTL2 and AD1/AD2 iPSC lines for all 3 methylation states across the genome, with 2 CTL iPSC lines with 4 replicates, and 2 AD iPSC cell lines with 4 replicates. (B) Violon plot of CG DMRs between CTL1/CTL2 and AD1/AD2 iPSC lines. Note the lower level of hyper-DMRs in ADs. (C) Venn diagram of DMR-associated genes between CTL1/CTL2 and AD1/AD2 iPSC lines for all 3 methylation states. (D) Gene ontology analysis of hyper and hypo DMR-associated genes between CTL1/CTL2 and AD1/AD2 iPSC lines for CG and CHH methylations. Note the predominance of DMR-associated genes linked to cell-cell adhesion, axonogenesis and synapses. (E) Venn diagram of DMR-associated transcription factors (TFs) between CTL1/CTL2 and AD1/AD2 iPSC lines for all 3 methylation states. (F) Top 9 DMR-associated transcription factors (TFs) between CTL1/CTL2 and AD1/AD2 iPSC lines for all 3 methylation states. (G) DMR-associated transcription factor (TF) families between CTL1/CTL2 and AD1/AD2 iPSC lines for all 3 methylation states.

### Reduced neural induction in AD neuroepithelial cells

Following exposure to BMP antagonists such as Noggin and using the “neural default model”, neural induction in human is first characterized by the expression of PAX6 and SOX1 in the primitive neural ectoderm ^69–71^. This stage is followed by the acquisition of neural progenitor cell identity, ultimately giving rise to all neuronal and glial cell types. We induced the differentiation of CTL and AD iPSCs into NE cells and performed RNA-seq analyses at DIV10 (**Fig. 3A**). PCA revealed that CTL and AD groups were relatively comparable, but with segregation of some samples (**Fig. S3A and B**). Heatmap and Volcano plot analyses showed numerous DEGs, with notable downregulation in ADs of genes involved in neural cell fate (NEUROD1, TBR1, UNCX, and LHX9), and upregulation of genes involved in cancer stem cell (ZIC2, ZIC5, and LGR4) and non-neuronal cell fate (ACTN3, VWF, and SERPINA5) (**Fig. 3B**). Interestingly, GSEA furthermore revealed activation of a stress response against Zinc and Cupper ion (**Fig. S3C**), similarly as observed in AD iPSCs (**Fig. 1I**). GO analysis revealed altered forebrain and telencephalon development in ADs (**Fig. 3C**). To further investigate this, we performed a developmental kinetics of CTL and AD NE cells from DIV5 to DIV20. By immunoblot analysis, we observed that in CTL NE cells, BMI1 and H2A^ub^ levels normally increase during the process of neural induction. In AD NE cells, BMI1 and H2A^ub^ levels were comparatively 30-50% lower at DIV15 and DIV20 (**Fig. 3D-E**). EZH2 levels were similar between CTL and AD NE cells at DIV15 but were reduced at DIV20 in AD NE cells (**Fig. 3D-E**). Notably, PAX6 and SOX1 levels were significantly lower in AD NE cells when compared to CTLs, suggesting less efficient neural induction (**Fig. 3D-E**). ZIC2 can activate WNT signaling ^72–74^, and LGR4 is a receptor for R-Spondin, which also activates WNT ^75^. To test the possible involvement of WNT, we exposed AD NE cells to XAV939 (a WNT inhibitor) or GSKJ1 (a histone demethylase inhibitor that promotes H3K27 bi- and tri-methylation) starting at DIV0. Notably, XAV939 treatment led to significant improvement of PAX6 and SOX1 levels and in reduction of apoptotic cells in ADs at DIV10, while positive effects of GSKJ1 treatment were more modest (**Fig. 3F-G**). *ZIC2* and *ZIC5* are co-regulated and are direct target of BMI1’s transcriptional repression activity ^76^. To investigate *BMI1* function in this context, we knockdown *BMI1* in NE cells starting at DIV0 (**Fig. 3H**). When compared to the siScramble control, we observed that *BMI1* knockdown in NE cells resulted in significant downregulation of PAX6 and SOX1, suggesting that *BMI1* is required for proper neural induction (**Fig. 3I-J**). Taken as a whole, these experiments revealed reduced neural induction and forebrain specification in AD NE cells, together with up-regulation of ZIC2/5 and of non-neuronal lineage genes.

**Figure 3.**
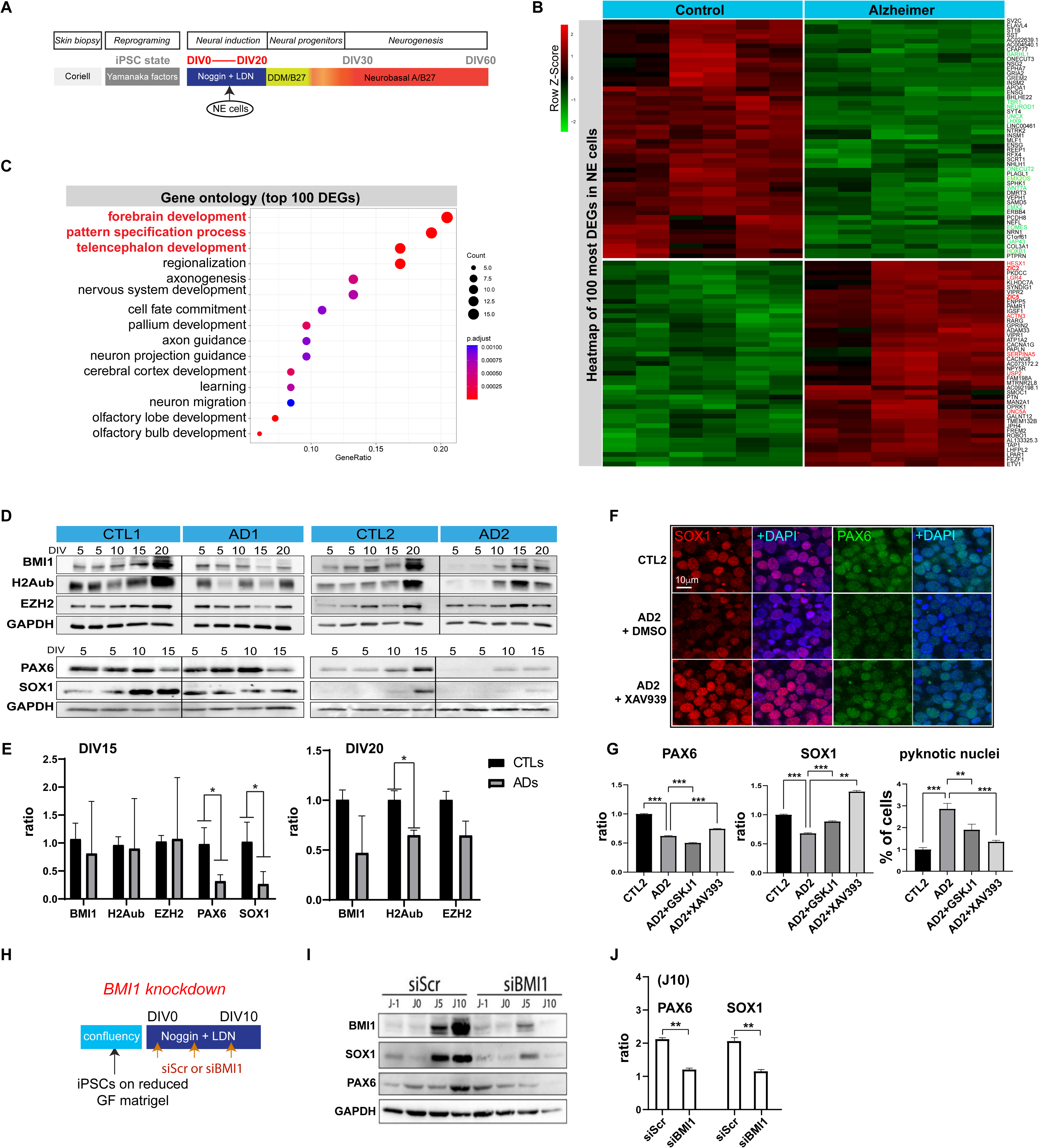
AD iPSCs show reduced neural induction efficiency. (A) Schematic of the cell’s differentiation steps used in this study to reach the neuroepithelial (NE) cell stage (DIV0-20). (B) Heatmap of unsupervised clustering of top 100 DEGs between CTL and AD NE cells at DIV10, with a cutoff of at least 2 folds. Genes of interest upregulated in ADs are in labeled in red. Genes of interest downregulated in ADs are in labeled in green. CTL = 3 cell lines with 6 replicates; AD = 2 cell lines with 4 replicates. (C) Gene ontology analysis of top 100 DEGs between CTL and AD NE cells at DIV10. (D) Western blot analysis of CTL and AD NE cells at DIV5 to DIV20. (E) Diagrams of quantification (n = 2 biological replicates). All values are mean ± SEM. (*) P< 0.05 by Student’s unpaired t-test. (F) Confocal immunofluorescence analysis of CTL and AD NE cells at DIV10, treated or not for 10 days with 10nM XAV939 or DMSO (n = 3 replicates). (G) Diagrams of quantifications, which also include data with 10nM of GSKJ1. (**) P < 0.01, (***) P < 0.001, SEM. Scale bar: 10mm. (H) Schematic of the method used for BMI1 knockdown in NE cells. (I) Western blot analysis of siScramble (siScr) and siBMI1 NE cells at J-1 (DIV-1) to J10 (DIV10). (J) Diagrams of quantification from (I) experiments at J10 (DIV10) (n = 3 replicates). All values are mean ± SEM. (**) P< 0.01 by Student’s unpaired t-test.

### WNT activation and mixed cell lineage identity in immature AD neurons

To follow the neural differentiation process, we analyzed CTL and AD cultures at DIV30 by IF (**Fig. 4A**). In contrast with CTL cultures showing diffuse p-Tau labeling in neurons, AD cultures presented p-Tau labeling in axons and within axonal swellings, as these were not present in MAP2-positive dendrites (**Fig. 4B**). Total Tau and F-actin immunoreactivity was also more intense in the AD1 culture (**Fig. S4A**) ^77^. Aβ42 accumulation was however not observed in either CTL or AD cultures. While the number of TuJ1-positive immature neurons was comparable between CTLs and ADs, the number of MAP2-positive neurons was slightly higher in ADs, suggesting possible accelerated neurogenesis (**Fig. 4B-C**). Quantitative analyses showed a ∼40% reduction in BMI1 and H2A^ub^ levels between CTLs and ADs, with variations also observed within CTL and AD groups (**Fig. 4C**). We next performed RNA-seq analysis of DIV30 cultures (**Fig. S4B**). Volcano plot analysis showed robust upregulation of *RSPO1, RSPO2 and RSPO3* in ADs (**Fig. 4D**). R-Spondins are secreted agonists of the WNT/β-catenin signaling pathway ^78^. Mining of existing databases also showed that 53% of up-regulated genes in AD cultures presented enrichment for PRC1 or PRC2 (**Fig. 4E**). Heatmap analysis of top 100 DEGs showed that downregulated genes in ADs were linked to neuronal maturation. On the other hand, many up-regulated genes in ADs were linked to mesoderm cell fate, including MSX1, MSX2, PAX3, PAX7, ZIC2, and ZIC5 (**Fig. 4F**) ^79–82,83^. Interestingly, both *ZIC2* and *MSX2* contain a Polycomb Repressive Element enriched for BMI1 ^76^. GO analysis further confirmed robust activation of the WNT pathway in ADs (**Fig. 4G**). We concluded that while AD cultures generate immature neurons in equal proportion as CTLs, they show non-neuronal lineage and WNT pathway gene expression.

**Figure 4.**
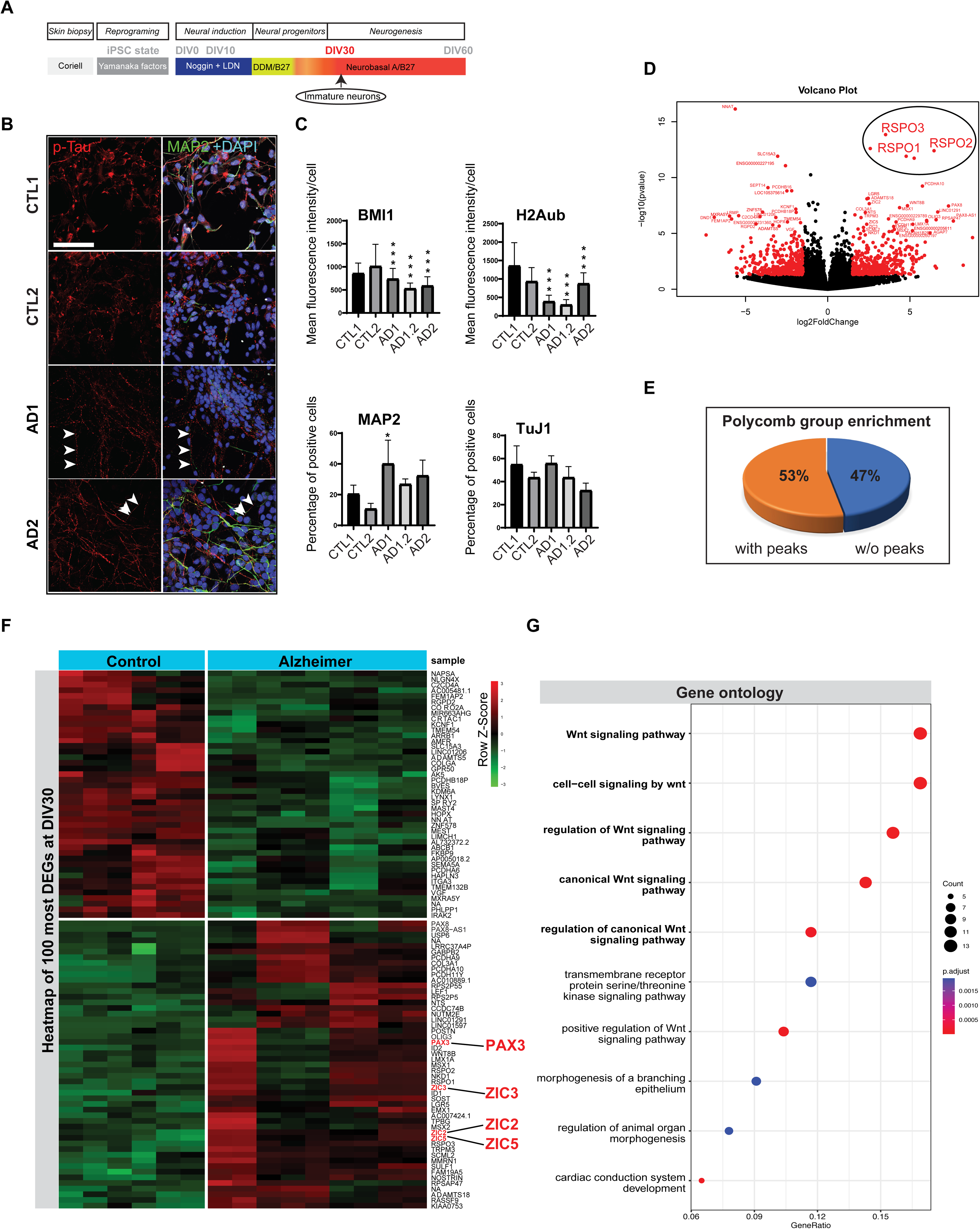
Immature AD neurons show mixed cell lineage identity and WNT activation. (A) Schematic of the cell’s differentiation steps used in this study to reach the immature neuronal cell stage (DIV30). (B) Confocal immunofluorescence analysis of CTL and AD neuronal cells at DIV30. White arrowheads point to p-Tau positive (in red) axonal swelling in AD neurons, and which are negative for the dendritic marker MAP2 (in green). (C) Diagrams showing quantification of confocal immunofluorescence analyses performed at DIV30 (n = 3 replicates). All values are mean ± SEM. (*) P< 0.05 and (**) P< 0.01 by Student’s unpaired t-test. (D) Volcano plot of all genes expressed in DIV30 neuronal cultures (CTLs vs ADs), with CTL = 3 cell lines with 6 replicates; AD = 4 cell lines with 8 replicates. Pluripotency genes are labelled in green. Significant DEGs at more than 2 folds (red dots). The RSPO1-3 gene family is overexpressed in AD cultures (circle). (E) Pie chart showing the proportion of upregulated DEGs in AD neuronal cultures at DIV30 and containing a Polycomb peak for either PRC1 or PRC2. (F) Heatmap of unsupervised clustering of top 100 DEGs between CTL and AD neuronal cultures at DIV30, with a cutoff of at least 2 folds. Genes of interest upregulated in ADs are in labeled in red. (G) Gene ontology analysis of all DEGs between CTL and AD neuronal cultures at DIV30, revealing significant activation of the Wnt signaling pathway in ADs.

### Dedifferentiation and loss of cell identity in AD neuronal cultures

We next analyzed CTL and AD neuronal cultures at DIV60 by IF to investigate late neuronal cell fate (**Fig. 5A**). In both CTL and AD cultures, ∼80% of the cells were positive for βIII-tubulin, and ∼60% positive for MAP2 (**Fig. 5B**). CTL and AD neurons also expressed markers of telencephalic origin, including FOXP2 and FOXG1, as well as GABAergic (GABA) and Cholinergic (ChAT) markers (**Fig. 5B**). The number of neurons per field was not significantly different between CTL and AD cultures (**Fig. 5B**). Bulk neuronal cultures were processed for RNA-seq analysis. Despite showing a net segregation and a large number of DEGs (**Fig. S5A and B**), CTL and AD cultures both expressed most neuronal sub-types present in the human cortex, including GABAergic, glutamatergic, Cajal-Retzius, upper layers and deep layers neurons (**Fig. 5C**). Notably however, AD cultures retained low expression of the pluripotency gene POU5F1 (**Fig. 5C**) and showed overall lower expression levels for neuronal genes (**Fig. 5C**). Deeper analysis confirmed higher expression of the POU5F1, SALL4, and LIN28A pluripotency genes, of the HOXB1, HOXB2 and HOXB3 polycomb target genes, and of the presomitic mesoderm and neural crest gene PAX3 in ADs (**Fig. 5D**). GO analysis confirmed that neurotransmitter transport, synaptic function, and neurotransmitter-gated ion channel activity were the most affected biological pathways in AD neuronal cultures (**Fig. 5E**). Moreover, GO analysis of major developmental pathways showed that generation of neurons was reduced in AD cultures, while BMP, WNT and TGF beta signaling pathways were all activated (**Fig. S5B**). To obtain a better understanding of this phenotype, we performed single-cell RNA-seq. UMAP of the major cell types found in DIV60 cultures showed the presence of Cholinergic, GABAergic and Glutamatergic neurons (**Fig. 5F**). Astrocytes, oligodendrocytes, radial glial cells, and quiescent neural stem cell-like cells (defined altogether here as glial cells) were also relatively abundant in the cultures (**Fig. 5F**). Comparative analysis of CTL and AD populations showed that glial cells in AD expressed higher levels of neural stem cell (*SOX2, NES,* and *NOTCH1*) and cell proliferation (*PCNA* and *MKI67*) markers (**Figs. 5G and S5D**). Notably, neurons in AD expressed lower levels of *MAP2*, while *MAP2* appeared ectopically expressed in glial cells (**Fig. 5H**). *CHAT*-expressing cells were notably less abundant in AD, either because of cell death, reduced genesis, or loss of cell identity (**Fig. 5H**). *PAX3* is expressed in the presomitic mesoderm and the neural crest ^82,82,84^. *PAX3* expression in glial cells and apparently also in neurons was elevated in AD cultures (**Fig. 5I**). While the fraction of MAP2 protein- and gene-expressing cells was comparable between CTL and AD (**Fig. 5J and K**), a larger fraction of cells in AD expressed *PAX3, PAX6, PAX7* and *VIM* (**Figs. 5K and S5C**). Looking only at *MAP2*-expressing cells, we found that *PAX3* and *VIM* were much more frequently co-expressed with *MAP2* in AD compared to CTL (**Fig. 5L**). Thus, neurons in AD express lower levels of neuronal genes and express genes of the mesoderm and neural crest lineage such as *PAX3*. In turn, glial cells in AD express higher levels of stem cell and cell proliferation-related genes and possibly also express neuronal genes.

**Figure 5.**
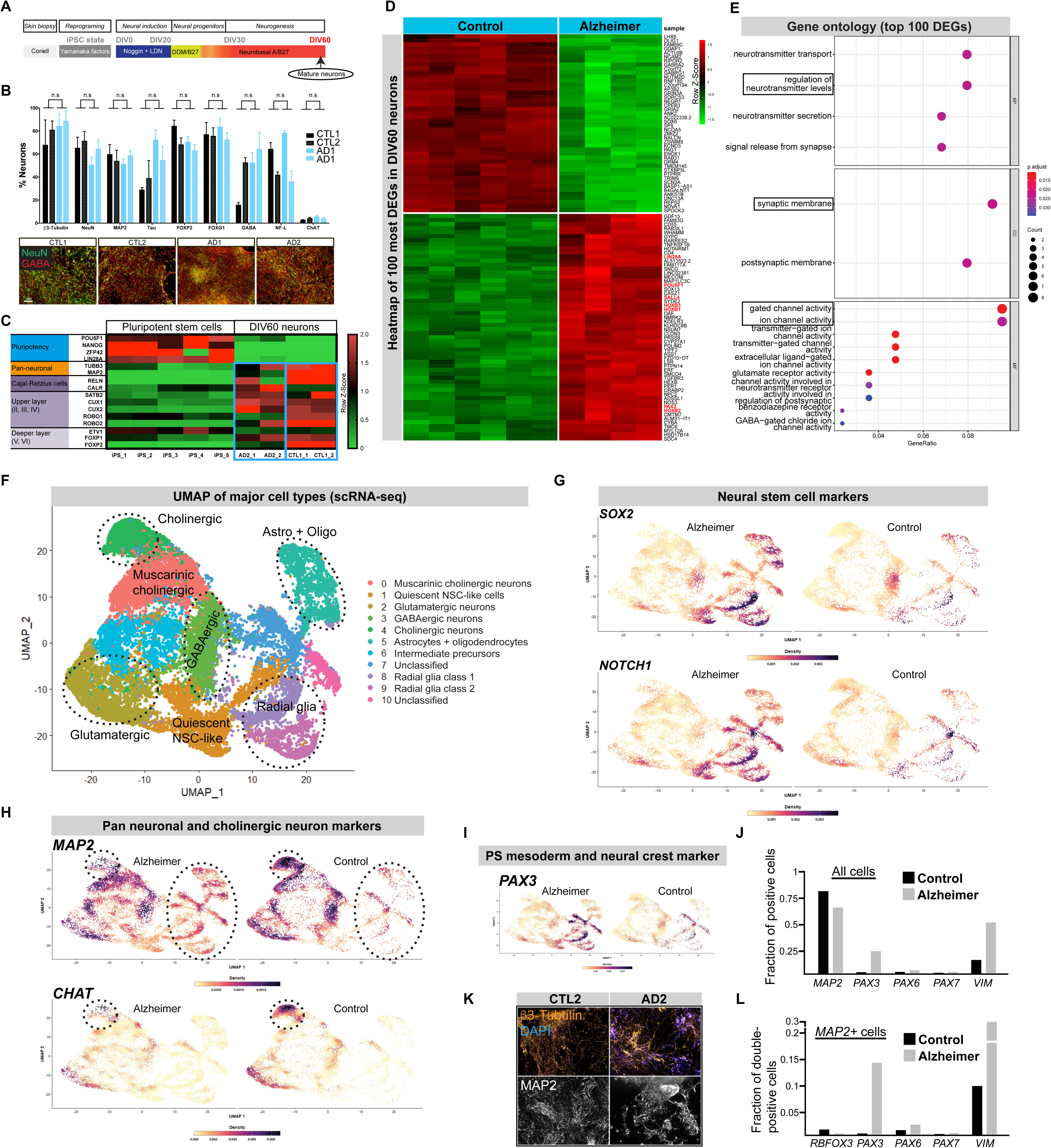
Dedifferentiation and loss of cell identity in AD neuronal cultures. (A) Schematic of the cell’s differentiation steps used in this study to reach the mature neuronal cell stage (DIV60). (B) Diagram of the percentage of positive neurons between CTL and AD neuronal cultures at DIV60 (n.s. = non-significant) as determined by confocal immunofluorescence (SEM, n = 3 replicates). Representative images of confocal immunofluorescence analyses of CTL and AD neuronal cultures at DIV60. Scale bar is 40mm. (C) Heatmap of selected pluripotency and neuronal genes between CTL and AD neuronal cultures at DIV60. Control induced pluripotent stem cells (iPS1-5) were used as blueprint of pluripotency. Note higher pluripotency gene expression, and lower neuronal gene expression in AD neuronal cultures at DIV60 when compared to CTLs at DIV60. (D) Heatmap of unsupervised clustering of top 100 DEGs between CTL and AD neuronal cells at DIV60, with a cutoff of at least 2 folds. CTL = 3 cell lines with 5 replicates; AD = 2 cell lines with 4 replicates. Selected genes overexpressed in ADs are in red. (E) Gene ontology analysis of all DEGs between CTL and AD neuronal cultures at DIV60, revealing significant alterations of neurotransmitter transport and regulation, synaptogenesis and ion channel activity in ADs. (F) Uniform Manifold Approximation and Projection (UMAP) of major cell types identified in CTL and AD neuronal cultures at DIV60 using scRNA-seq analysis. (G) UMAP plot comparison between CTL and AD neuronal cultures at DIV60 showing upregulation of neural stem cell markers in AD cells. (H) UMAP plot comparison between CTL and AD neuronal cultures at DIV60 showing ectopic expression of the pan neuronal marker MAP2 in AD glial cells (dotted circles) and reduced *MAP2* and *CHAT* expression in AD neurons (dotted circles). (I) UMAP plot comparison between CTL and AD neuronal cultures at DIV60 showing increased and ectopic expression of the *PAX3* in AD cells. (J) Diagram showing the fraction of positive cells for various neuronal and non-neuronal markers in CTL and AD cultures at DIV60. (K) Confocal immunofluorescence analyses of CTL and AD neuronal cultures at DIV60 showing comparable numbers of b3-Tubulin and MAP2-positve neurons. Scale bar: 40mm. (L) Diagram showing the fraction of MAP2 double-positive cells for various neuronal and non-neuronal markers in CTL and AD cultures at DIV60. Note the elevated proportion of *MAP2*-positive neurons also expressing *PAX3* or *VIM* in AD cultures.

### Mitigation of pathology in AD neuronal cultures by BMI1 overexpression

To test if AD iPSCs could differentiate into organoids despite their epigenomic anomalies, we generated floating brain organoids ^85^. Organoids measuring ∼1-2 mm in diameter were obtained from CTL1, CTL2, AD1 and AD2 iPSC lines and harvested at DIV90 (**Figs. 6A and S6A**). Both CTL and AD brain organoids frequently showed an internal cavity and expressed MAP2 (**Fig. 6A**). In contrast with CTLs however, AD brain organoids were immunoreactive for Aβ42 using 3 distinct antibodies, revealing intra- and extra-cellular accumulation of Aβ42-positive fiber-like structures (**Fig. 6A and A’**). Amyloid plaques were not observed. AD brain organoids also showed p-Tau accumulation, which was not present in CTL organoids (**Fig. S6B**). Likewise, AD1 and AD2 neuronal cultures at DIV60 showed many anomalies as revealed using immunoblot. When compared to CTLs, total APP level was higher in AD1 cultures, while level of the C88/C99 fragment was higher in AD2 cultures (**Fig. 6B**). These marked pathological differences were also observed when looking at BACE1, GSK3β, p-GSK3β (Ser9) and total Tau levels (**Fig. 6B**). Swelling of dendrites, Aβ42 accumulation, and elevated Aβ42 secretion was observed in AD3 cultures, as previously reported, together with reduced BMI1 levels (**Fig. S6C-F**) ^34,45^. Reduced BMI1, H2A^ub^, and H3K9^me3^ levels were also observed in AD3 cultures using immunoblot (**Fig. S6G**), suggesting loss of heterochromatin. Hence, AD1 and AD2 neurons presented severe anomalies in the expression and distribution of H3K9^me3^ and LaminA/C, as revealed by IF (**Fig. 6C and D**). In some cases, the nuclear envelope showed a “butterfly-like” morphology reminiscent of laminopathies (**Fig. 6C**) ^86^. To test if *BMI1* knockout could perturb the nuclear envelope, we inactivated *BMI1* using CRISPR/Cas9 in DIV50 neuronal cultures produced from the LMNB1-GFP iPSC line. When observed at DIV60, sgRNA-CTL neurons presented a robust and relatively uniform LMNB1-GFP and H3K9^me3^ signal (**Fig. 6E**). In contrast, sgRNA-BMI1 neurons showed reduced H3K9^me3^ staining and nuclear envelope anomalies (**Fig. 6E**). BMI1 overexpression can reduce p-GSK3β and p-P53 levels in cultured human neurons and is protective against camptothecin or 3-NP toxicity in mouse neurons ^34,87^. The PRC1 also promotes chromatin compaction and higher order chromatin structure ^88,89^. To test if BMI1 overexpression could reverse the observed AD’s neuronal pathology, we used lentiviral infection or multiple plasmid transfection (**Fig. 6F**). Lentiviral infection effectively increased BMI1 expression in AD cultures (**Fig. 6G**). It also reduced p-Tau and Aβ42 levels, and secreted Aβ42 concentration (**Figs. 6H-I**). Chromatin compaction is required to prevent the formation of G-quadruplex DNA structures (G4s), and *BMI1* inactivation in neurons results in the induction of G4s ^90^. Consistently, we observed that G4s in AD neurons were associated with increased histone H3K9 acetylation, a mark of relax chromatin (**Fig. S6H**) ^91^. Notably, repeated transfection of the BMI1 plasmid starting at DIV30 or DIV50 significantly reduced the number of G4-positive cells in AD cultures at DIV45 and DIV60, respectively (**Fig. 6J and K**). When compared to control-plasmid transfected AD cells, BMI1 overexpression in AD cells at DIV30 also promoted the formation of a ring-like H3K9^me3^-labeled heterochromatin at DIV45 (**Fig. 6J**). BMI1 overexpression in AD cells at DIV50 also significantly reduced heterochromatin fragmentation at DIV60 (**Fig. 6K**). These experiments suggest that BMI1 overexpression can mitigates multiple phenotypes observed in AD neuronal cultures.

**Figure 6.**
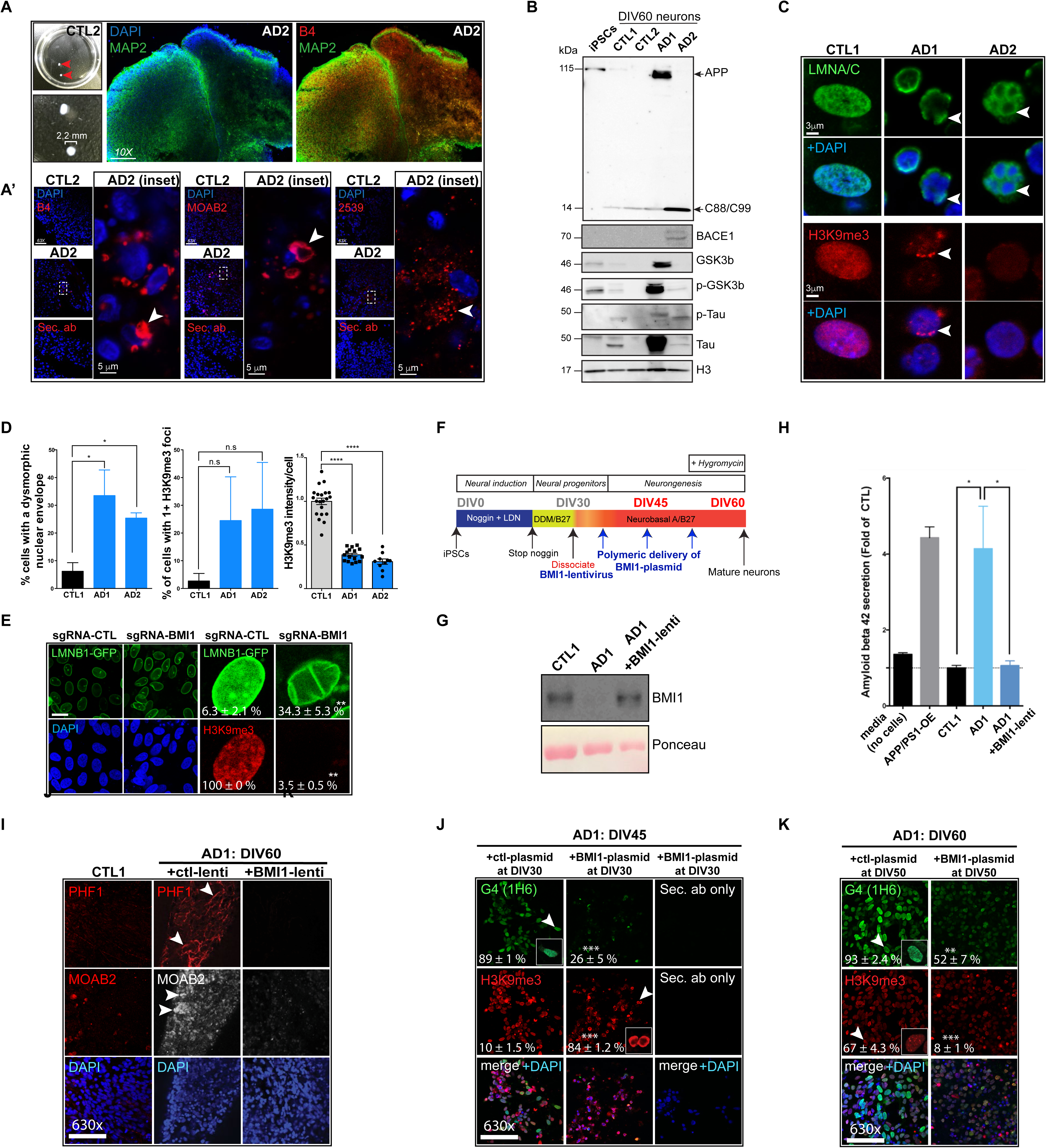
Neuronal pathologies in AD neurons are mitigated by BMI1 overexpression. (B) Left: Images of brain organoids formed using CTL1 iPSC lines after 3 months of maturation. Right: Dendritic staining (MAP2) and ý-amyloid (B4) staining in cryosections from AD2 brain organoids after 3 months of maturation. Scale bar: 100 mm. (A’) Extra-cellular ý-amyloid accumulation in AD2 brain organoids but not in CTL1 brain organoids after 4 months of differentiation using 3 independent antibodies (B4, MOAB2 and ab2539). The white arrow shows extra-cellular deposits. AD2 brain organoids stained only with the secondary antibody was used to assay for possible non-specific background fluorescence. Scale bar: 5mm. (C) Immunoblot on CTL1, CTL2, AD1 and AD2 neuronal cultures for APP, BACE1, GSK3beta, p-GSK3beta (Ser9), phospho-Tau (PH13) and Tau (K9JA). H3 was used as loading control. Undifferentiated iPSCs were used as a negative control. (D) Immunostaining for LMNA/C and H3K9me3 in CTL1, AD1 and AD2 neuronal cultures after 2 months of differentiation. Note the abnormal nuclear envelope stained with LMNA/C and the fragmentation of the heterochromatin in the AD neurons (arrowheads). Scale bar: 3mm. (E) Quantification of the number of dysmorphic nuclear envelopes and H3K9me3 foci in CTL1, AD1 and AD2 neuronal cultures (n = 4 for each culture). All values are mean ± SEM. (*) P< 0.05 by Student’s unpaired t-test. (F) Confocal immunofluorescence of control (sgRNA-CTL) and BMI1 knockout (sgRNA-BMI1) neuronal cultures produced from the differentiation of the LMNB1-GFP iPSC line. Note the laminopathy and loss of heterochromatin following BMI1 knockout. All values are mean ± SEM. (**) P< 0.01 by Student’s unpaired t-test. Scale bar: 3mm. (G) Scheme depicting the method and timeline of differentiation for overexpressing BMI1 at the neural progenitor stage or at the time of neurogenesis. (H) Immunoblot on CTL1, AD1, and AD1 neuronal cultures infected with the BMI1-expressing lentivirus. Ponceau was used as loading control. (I) Extracellular Aβ342 levels in CTL1, AD1, and AD1 neuronal cultures infected with the BMI1 lentivirus (n = 16; 2 lines; 4 biological replicates in 2 technical replicates) as measured by ELISA. Neurons overexpressing APP- and PSEN1-mutant transgenes were used as positive control. Media not conditioned is used as negative control. All values are mean ± SEM. (*) P< 0.05 by Student’s unpaired t-test. (J) Immunostaining of DIV60 CTL and AD neuronal cultures. Accumulation of p-Tau and β-amyloid in AD1 neurons is indicated (arrowheads). Note the absence of staining in AD1 cultures expressing the BMI1 lentivirus. Scale bar: 40mm. (K) Immunostaining for G4 DNA using the 1H6 antibody (arrowhead-inset) and H3K9me3 in AD1 with or without BMI1 overexpression after 45 days of differentiation. Note the ring-like structure of H3K9me3 staining upon BMI1 over-expression (arrowhead-inset). Numbers indicate the number of positive cells for each marker (n = 3 independent images). All values are mean ± SEM. (***) P< 0.001 by Student’s unpaired t-test. (L) Immunostaining for G4 DNA using the 1H6 antibody (arrowhead-inset) and H3K9me3 in AD1 with or without BMI1 overexpression after 60 days of differentiation. Note heterochromatin fragmentation labeled with H3K9me3 in the ctl-plasmid AD1 cultures (arrowhead-inset). Numbers indicate the number of positive cells for each marker (n = 3 independent images). Heterochromatin fragmentation was defined by five or more H3K9^me^^3^-positive chromocenters/nucleus. All values are mean ± SEM. (**) P< 0.01 and (***) P< 0.001 by Student’s unpaired t-test.

## DISCUSSION

Herein, we have showed that AD iPSCs, while being pluripotent, retain epigenomic anomalies characterized by reduce *BMI1* expression, lower H3K9^me3^ levels, and altered DNAme. GO terms associated with AD iPSC’s DMRs were reminiscent of the Alzheimer’s resilient epigenetic signature, with MEF2s enrichment at DMRs. Reduced neural induction and forebrain specification in AD NE cells was associated with upregulation of ZIC2, ZIC5, LRG4 and of non-neuronal genes. Robust WNT signaling and expression of mesoderm cell fate genes was also observed at DIV30. This culminated in de-repression of cell proliferation and stem cell genes at DIV60, together with apparent loss of cell identity and depletion of cholinergic neurons. At both DIV30 and DIV60, however, we observed that the number of AD neurons produced was similar to that of the control neurons. Likewise, AD iPSCs were able to differentiate into more complex structures such as brain organoids. Notably, many of the pathologies observed in AD neurons were mitigated following BMI1 overexpression.

Following differentiation of iPSCs toward neural ectoderm, AD neurons and glia cells presented a phenotype of de-differentiation and loss of cell identity. A comparable phenomenon was described in AD neurons and glial cells *in situ*, as revealed using single-nuclei RNA-seq of late AD brains and referred to as epigenome erosion ^14^. Similarly, induced neurons from AD HDFs show reduced neuronal maturation, activation of stress response genes, and increased expression of stem cell and cancer-related genes ^47^. While the underlying mechanism of these observations are unknown, it is plausible to stipulate that de-differentiation cause by reactivation of developmental genes is sufficient to explain this phenotype. In human pluripotent stem cells, the non-canonical PRC1 is composed of PCGF6 and CBX7. Upon neural induction, these are replaced by the canonical PRC1 proteins BMI1 and CBX8. PCGF6 is enriched at WNT-related genes and *PCGF6*-null iPSCs show upregulation of meso-endoderm genes and downregulation of neuronal genes ^92^. Similarly, *Phc1*-null mouse embryonic stem cells fail to differentiate into neural tissue and show upregulation of pluripotency and non-neural genes ^93^. Here we found that reduced neural induction in AD iPSCs correlated with lower BMI1 and H2A^ub^ levels and activation of *ZIC2*, ZIC5 and *LRG4*. Neural induction in AD NE cells was improved following WNT inhibition. Conversely, *BMI1* knockdown impaired neural induction in NE cells. It is notable that the *ZIC2* gene contains a Polycomb Response Element at its promoter and is a direct target of BMI1 repressive activity ^76^. *ZIC2* and *ZIC5* are coregulated and can activate WNT and TGF beta signaling as well as *POU5F1* transcription to promote a cancer stem cell phenotype ^72,73,94–96^. Thus, lower PRC1 activity in AD iPSCs could explain activation of *ZIC2*, *ZIC5* and of WNT signaling. During WNT signaling, β-catenin interacts with TCF and recruits’ transcriptional co-activators and histone modifiers, and TCF binding sites are found in *SALL4, SOX2, NANOG*, and *POU5F1* (The WNT Homepage: https://web.stanford.edu/group/nusselab/cgi-bin/wnt/). Thus, sustained WNT signaling could explain reactivation of pluripotency, cell proliferation and cancer-related genes in AD cultures.

It has been estimated that, when compared to human embryonic stem cells, 60-85% of DMRs in primed iPSCs originate from the cell of origin ^42,43^. Work on iPSCs and nuclear transfer embryos also revealed that these reprogramming resistant chromatin regions are enriched for H3K9^me3^ ^42,44^. Here we found that the H3K9^me3^ histone mark is the most affected in AD iPSCs, and that AD iPSCs present DMRs in genes associated with cell-cell adhesion and synapse, with robust enrichment for the MEF2 transcription factors. MEF2s are key genes in cognitive resilience and are required for normal neurodevelopment and synaptogenesis ^9,58,59,63,97^. Thus, abnormal DNAme of MEF2s target genes in AD iPSCs is predicted to perturb transcription at the time of neuronal differentiation. This could explain other anomalies observed in AD cultures such as those affecting synaptic membrane, neurotransmitter secretion, and neurotransmitter transport (**Fig. 5E**). While the AD pathology is largely restricted to the brain, it is intriguing that HDFs can transmit the disease imprint after reprogramming into iPSCs, and also following neuronal differentiation. Remaining within the iPSC experimental paradigm, these findings suggest either the presence of uncharacterized genetic alterations working in trans on the epigenome, or the presence of AD-specific epigenomic anomalies that are reprogramming resistant. One way to decipher between genetic and epigenomic would be to perform transient-native-treatment reprogramming of AD HDFs into iPSCs ^42^. This method erases most reprogramming resistant regions, producing iPSCs with an epigenome that is near identical to that of human embryonic stem cells. Rescue of the epigenomic anomalies observed in AD iPSCs would support an epigenomic-origin, while maintenance of the anomalies would support a genetic-origin.

In conclusion, we have demonstrated that the epigenome of AD iPSCs differs from that of normal control iPSCs, and that these epigenomic anomalies relate to pathways highly relevant to AD. Likewise, neural differentiation of AD iPSCs recapitulated several phenotypes observed in AD brains, suggesting that the epigenomic alterations are retained during differentiation. In depth characterization of the AD epigenome could have broader implication in our understanding of the disease origin and pathophysiological mechanisms, and possibly holds promises for the development of new therapeutics targeting the epigenome.

### Limitations of the study

Neuronal cultures and brain organoids from AD iPSCs cannot reflect the complexity present in a living organism, lacking immune cells, myelination, and blood circulation amongst other things. AD is also an adult-onset neurodegenerative disease not affecting development, while the iPSC model recapitulates development before generating mature neuronal cells. In this study, all AD iPSC lines were males. Female AD iPSCs may have revealed distinct epigenome modifications and neuronal pathologies that are not present in male AD iPSCs. Comparative DNA methylome analysis was not performed between donor’s fibroblasts and reprogramed iPSCs.

## ACKNOWLEDGMENTS

This work was supported by grants from the National Science and Engineering Research Council of Canada (NSERC), Maisonneuve-Rosemont Hospital Foundation, and Pierre Théroux Family Foundation for Alzheimer’s Disease Research. Rimi Hamam was supported by a fellowship from Maisonneuve-Rosemont Hospital Foundation. We thank Dr. Patrick Nkanza for histological analysis of teratomas.

## Data availability statement

Raw data, cell lines, and reagents are available upon request.

## Code availability

To review GEO accession GSE255985; SubSeries GSE255981, GSE255982, and GSE255983: Go to https://www.ncbi.nlm.nih.gov/geo/query/acc.cgi?acc=GSE255985 Enter token khyhokoibpqfjml into the boxSeries GSE255985

## Author contributions

Conceived and designed: G.B., AK, RH, and AF. Performed the experiments: AF, AK, DS, RH, RyH, and RiH. Analyzed the data: GB, AF, ET, and RH Wrote the paper: GB, AF, and RH

## MATERIALS AND METHODS

### Human samples

Human pluripotent stem cells were approved by the “Comité de Surveillance de la Recherche sur les Cellules Souches” of the CIHR and Maisonneuve-Rosemont Hospital Ethic Committee and used by following the Canadian Institute Health Research (CIHR) guidelines. All methods were carried out in accordance with relevant guidelines and regulations. Controls and AD fibroblasts were obtained from clinically diagnosed individuals from Coriell Biorepository, with informed consent from all participants.

### Generation of human induced pluripotent stem cells

Human dermal fibroblasts were ordered from the Coriell Institute. We selected an 82-year-old male and a 92-year-old female with no cognitive impairments as elderly controls (CTL1: #AG04152, and CTL2: #AG09602, respectively). We selected a 72-year-old male deceased with sporadic late-onset AD (AD1: #AG08243) and with no familial history of AD, and a 90-year-old male (AD2: #AG08259) also deceased with sporadic late-onset AD and with no familial history of AD. This later patient had dementia since the age of 87 years. The presence of p-Tau tangles and amyloid plaques in the brains of both patients was confirmed in post-mortem studies. Other CTL and AD samples are described in Figure 1C. The reprogramming of dermal fibroblasts was based on a previous method ^49^. Briefly, dermal fibroblasts were expanded (using DMEM/F12 + 10% FBS media) and transfected by 3 plasmids for the expression of Yamanaka reprogramming factors (pCXLE-hSK, pCXLE-hOCT3/4-shp53-F and pCXLE-hUL). After 7 days, transfected cells were transferred onto irradiated Mouse Embryonic Fibroblasts (MEFs) and further cultured in ReproTeSR media (StemCell Technologies). After 25-30 days of transfection, iPSC clones were manually picked, expanded, and characterized.

### Differentiation of iPSCs into cortical neurons and cortical brain organoids

The differentiation protocol was based on a previous study. However, the Noggin agonist LDN193189 was used to reduce recombinant Noggin concentration. The CTL1, CTL2, sAD1 and sAD2 cell lines were dissociated using Accutase (Innovative Cell Technology #AT-104) and platted on growth factor reduced Matrigel (Corning #356231) in PeproGrow hES cell media (PeproTech #BM-hESC) supplemented with ROCK inhibitor (Y-27632;10µM, Cayman Chemical #10005583). Upon 70% of confluency, the media was changed to DDM supplemented with B27 (1X final), Noggin (10 ng/ml, PeproTech #120-10C) and LDN193189 (0.5µM; Sigma #SML0559) resulting in the DDM/B27/LN medium. The medium was changed every day. After 16 days of differentiation, the medium was changed to DDM/B27 and replenished every day. At day 24, neural progenitors were manually detached from the plate and platted on growth factor reduced matrigel coated plates or chamber slides (LabTek #154534). Five days after the dissociation/infection, half of the medium was changed for Neurobasal A media supplemented with B27 (1X final) and changed again every three days. For cortical brain organoids, iPSCs were induced to produce embryonic bodies by adding 5% serum in DDM/B27 and using ultra-low cell culture plates. After 7 days, embryonic bodies were coated with Matrigel and cultured in serum-free DDM/B27/LN medium until DIV30. At this point, brain organoids were maintained in DDM/B27 medium and replenished every 3 days.

### Bulk RNA-seq

Total RNA from CTL and AD samples were extracted iPSC, iPSC differentiated to neurons for 10, 30 and 60 days, using the standard procedure of Qiagen columns and assayed for RNA integrity. cDNA was prepared according to the manufacturer’s instructions (NEB library) and sequenced using the Illumina platform. Base-calling and feature count were done using Illumina software. For differential expression analysis, DESeq2 ^98^ was used on R program (Team, R. C. The R Project for Statistical Computing. Http://Www.R-Project.Org*<u>/</u>* 1–12 (2013) at https://www.r-project.org/). Global gene expression was visualized using volcano plots and heatmaps for top genes. Samples were clustered and visualized in a 2D space using PCA ^99^. For functional analysis, the differentially expressed genes, or a subset of those genes, were analyzed using Gene Ontology (GO), Gene Set Enrichment Analysis (GSEA) and Reactome ^100–105^.

### bWGS

Total DNA from CTL and AD iPSC cell lines were extracted using Qiagen Kit according to manufacturer’s instructions and processed for a whole genome bisulfate sequencing. To obtain the qualified clean data, we trim the raw reads to filter out the contaminated adapter sequence and low-quality reads using Trimmomatic. Bismark software was used to perform alignments of bisulfite-treated reads to the HG38 reference genome and to identify differentially methylated regions (DMRs) ^106^. The criteria for calling a DMR were as the sequencing depth is greater than or equal to five; the q-value less than or equal to 0.01. DMRs were then annotated for the genes that are affected by them, these genes were compared using a Venn Diagram to establish the genes that are affected by the CG, the CHG, the CHH modifications or by any combination of these modifications. We performed a GO analysis on the genes that contain a DMR, as well as a KEGG analysis to establish the functions of these genes and the pathways affected by these differential methylations. The DMRs were analyzed for their affinity to transcription factors using the JASPAR software, and methylation levels were visualized using the IGV software ^100,101,106–108^.

### Single-cell RNA-seq analysis

Cells were dissociated and prepared with the standard procedure of 10X Single-cell gene expression kit aiming for the collection of 10,000 cells, with 1 AD sample and 1 CTL sample. The resulting library was sequenced with the Illumina platform. Base-calling and feature count were done using Illumina software. Read count was performed with Cell Ranger software 3.0.0 from 10X to analyze and map the reads on the HG38 genome ^109^. Loupe 4.1.0 was instrumental in visualizing the data and performing an unbiased ontology analysis to identify the cell lineage present in our cultures ^110^. In summary, cells were clustered without bias using the K-mean strategy on the UMAP distribution ^111^. The differentially expressed genes in each group identified by the cell ranger were compared to the literature and their ontology origins to identify the corresponding group. Using a curated gene list, we were able to assign to each group a cell type, adding the Seurat package for further analysis on these groups and for gene expression visualization ^112–114^.

### ELISA

ELISA assays (Invitrogen #PP0812) were done on cellular supernatants. Cell supernatants were sonicated before the assay. Extracts were processed according to manufacturer’s instructions. Cellular extracts were used to normalize to the total quantity of proteins.

### Immunofluorescence microscopy

All secondary antibodies were tested alone or in combination to assay for possible non-specific background fluorescence. Cells were fixed with 4% PFA for 15 min and permeabilized with Triton X-100 for 10 min. Unspecific antigen blocking was performed using 1% BSA in PBST for 30 min. Cells were incubated with the primary antibody overnight at 4C in a humidified chamber. After incubation with the secondary antibody, slides were counter stained with DAPI. Pictures were taken using a confocal microscopy system (Olympus). Brain organoids were fixed overnight with 4% PFA, dehydrated using sucrose gradient baths and cryo-sectioned (20μm) prior to permeabilization.

### Antibodies list

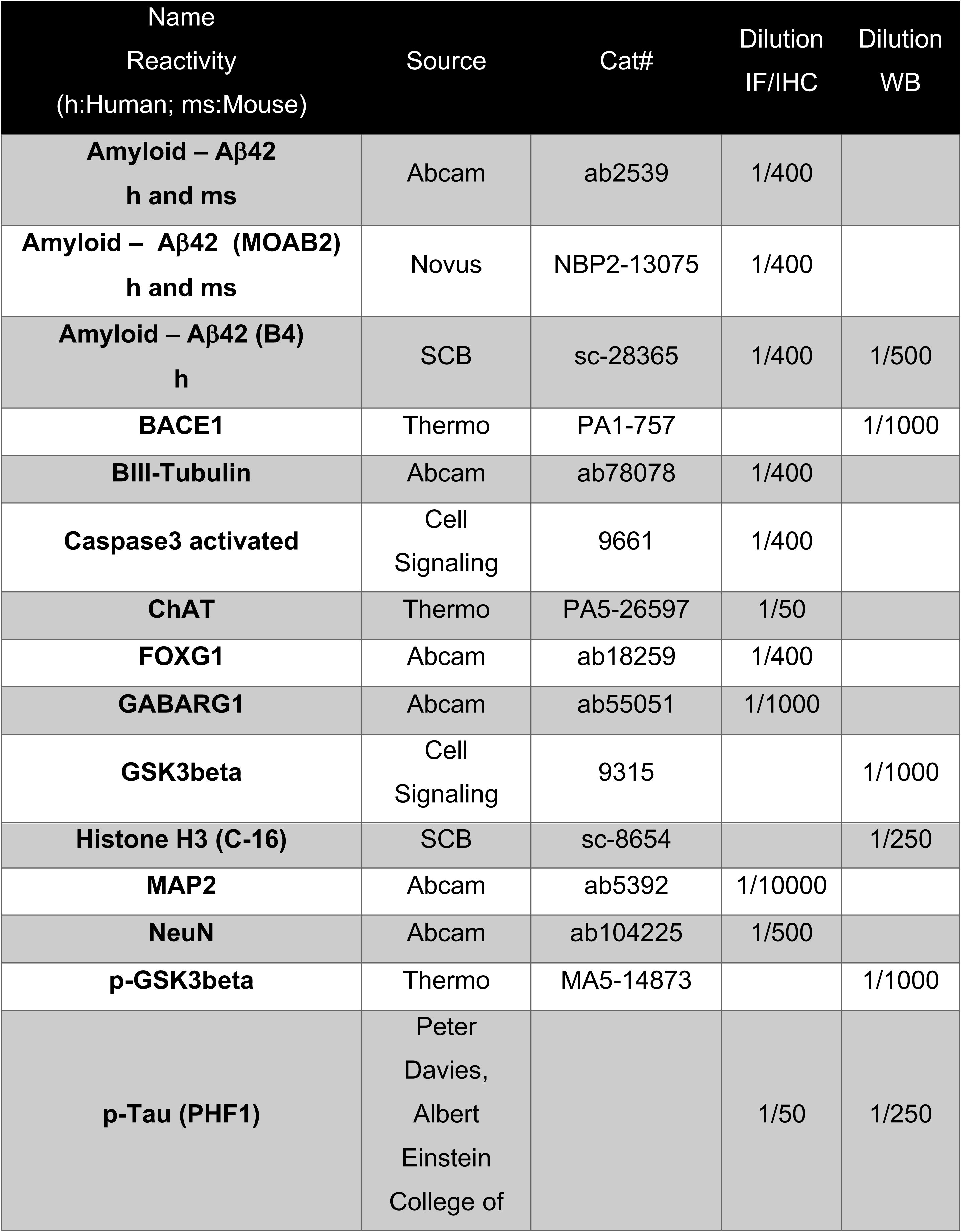
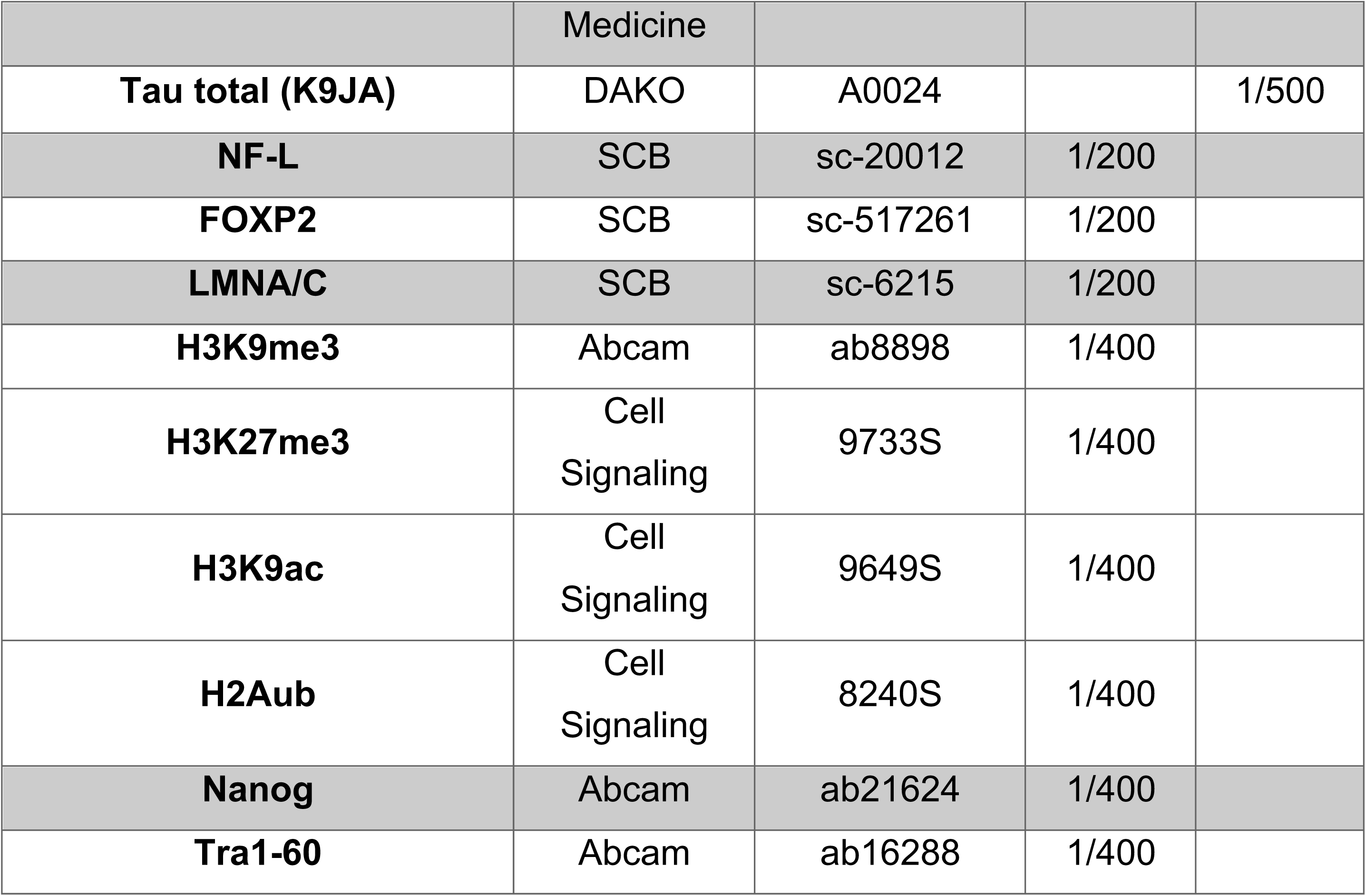

**Antibodies list:**
| **Name**<br>**Reactivity**<br>**(h:Human; ms:Mouse)** | **Source** | **Cat#** | **Dilution**<br>**IF/IHC** | **Dilution**<br>**WB** |
| --- | --- | --- | --- | --- |
| **Amyloid – Aβ42**<br>**h and ms** | Abcam | ab2539 | 1/400 |  |
| **Amyloid – Aβ42 (MOAB2)**<br>**h and ms** | Novus | NBP2-13075 | 1/400 |  |
| **Amyloid – Aβ42 (B4)**<br>**h** | SCB | sc-28365 | 1/400 | 1/500 |
| **BACE1** | Thermo | PA1-757 |  | 1/1000 |
| **BIII-Tubulin** | Abcam | ab78078 | 1/400 |  |
| **Caspase3 activated** | Cell<br>Signaling | 9661 | 1/400 |  |
| **ChAT** | Thermo | PA5-26597 | 1/50 |  |
| **FOXG1** | Abcam | ab18259 | 1/400 |  |
| **GABARG1** | Abcam | ab55051 | 1/1000 |  |
| **GSK3beta** | Cell<br>Signaling | 9315 |  | 1/1000 |
| **Histone H3 (C-16)** | SCB | sc-8654 |  | 1/250 |
| **MAP2** | Abcam | ab5392 | 1/10000 |  |
| **NeuN** | Abcam | ab104225 | 1/500 |  |
| **p-GSK3beta** | Thermo | MA5-14873 |  | 1/1000 |
| **p-Tau (PHF1)** | Peter<br>Davies,<br>Albert<br>Einstein<br>College of |  | 1/50 | 1/250 |

|  | Medicine |  |  |  |
| --- | --- | --- | --- | --- |
| **Tau total (K9JA)** | DAKO | A0024 |  | 1/500 |
| **NF-L** | SCB | sc-20012 | 1/200 |  |
| **FOXP2** | SCB | sc-517261 | 1/200 |  |
| **LMNA/C** | SCB | sc-6215 | 1/200 |  |
| **H3K9me3** | Abcam | ab8898 | 1/400 |  |
| **H3K27me3** | Cell<br>Signaling | 9733S | 1/400 |  |
| **H3K9ac** | Cell<br>Signaling | 9649S | 1/400 |  |
| **H2Aub** | Cell<br>Signaling | 8240S | 1/400 |  |
| **Nanog** | Abcam | ab21624 | 1/400 |  |
| **Tra1-60** | Abcam | ab16288 | 1/400 |  |

### Quantification of immunofluorescence images

IMARIS station v8.4.1 (Bitplane) was used for all quantification, using a Scholl regression coefficient was set at 1μm.

### Western Blot

Cell extracts were homogenized in the Complete Mini Protease inhibitor cocktail solution (Roche Diagnostics), followed by sonication. Protein material was quantified using the Bradford reagent. Proteins were resolved in 1x Laemelli reducing buffer by SDS-PAGE electrophoresis and transferred to a Nitrocellulose blotting membrane (Bio-Rad). Subsequently, membranes were blocked for 1h in 5% non-fat milk-1X TBS solution and incubated overnight with primary antibodies. Membranes were then washed 3 times in 1X TBS; 0.05% Tween solution and incubated for 1h with corresponding horseradish peroxidase-conjugated secondary antibodies. Membranes were developed using the Immobilon Western (Millipore). Blots were quantified using the Image quant program.

### Real-time RT-PCR

RNA was isolated using TRIzol reagent (Invitrogen). Reverse transcription (RT) was performed using 1 µg of total RNA and the MML-V reverse transcriptase (Invitrogen). Real-time PCR was carried in triplicates using Platinum SYBRGreen Supermix (Invitrogen) and Real-time PCR apparatus (ABI prism 7002). Primers used are:

ZFP42_F AGAAACGGGCAAAGACAAGAC
ZFP42_R GCTGACAGGTTCTATTTCCGC
NANOG_F AAGGTCCCGGTCAAGAAACAG
NANOG_R CTTCTGCGTCACACCATTGC POU5F1_F CTTGAATCCCGAATGGAAAGGG
POU5F1_R GTGTATATCCCAGGGTGATCCTC

### Detection of potential plasmid integration

Genomic DNA was extracted using DNA/RNAeasy kit from Qiagen from each iPSC line and then subjected to real-time PCR as previously described ^49^. Fibroblasts transfected with the 3 plasmids and collected at day 6 were used as positive control.

### Teratoma formation

2×10^5^ iPSCs from each line were collected and re-suspended into DMEM/F12/1%Matrigel supplemented with ROCK inhibitor (Y-27632;10µM, Cayman Chemical #10005583). Each line was injected into 2 flanks of 6-week-old NOD/SCID mice. When the tumor was greater than 0,5 cm the teratoma was extracted and included into paraffin blocks for histology.

### Statistical analysis

Statistical analysis was performed using Graphpad software (Prism 6). Statistical differences were analyzed using Student’s *t*-test for unpaired samples. In all cases, the criterion for significance (*P* value) was set as mentioned in the figures. For DMRs, criterion for significance was that the sequencing depth is greater than or equal to five; (2) q-value less than or equal to 0.01. When comparisons were made using independent samples of equal size and variance following a normal distribution, significance was assessed using an unpaired two-sided Student’s t-test. Where several groups were compared, significance was assessed by ANOVA and adjusted for multiple comparisons using the Bonferroni correction.

### Whole Genome Sequencing

See Supplementary Method

