## Supplementary material for "Epigenomic anomalies in induced pluripotent stem cells from Alzheimer’s disease cases": Methods, Figures and Tables

[7] Present address: McGill University, Montréal, Canada

* Senior author

**Corresponding author

### SUPLEMENTARY METHOD

#### Whole Genome Sequencing

Paired-end (2 x 150 bp) Whole Genome Sequencing (WGS) was performed using Illumina Hi-Seq X with mean coverage of 39.4 X for three affected (AD1, AD2, and FAD7) and three control samples (CTL1, CTL2, and CTL14), and Illumina NovaSeq 6003 with mean coverage 38.7% for two affected samples (FAD24, FAD25) and one control sample (GB16). Base calling for the samples run in Illumina HiSeqX was performed using Illumina HiSeq Analysis Software (HAS; version 2-2.5.55.1311). Reads were mapped to the human b37 reference sequence using bwa-mem v0.7.12 [1]. Duplicate reads were removed using MarkDuplicates from Picard v2.5.0. Local read realignment around indels, base quality score recalibration (BQSR), variant calling with HaplotypeCaller, and variant quality score recalibration (VQSR) were accomplished using GATK v3.7.0 [2]. Illumina Dragen DNA platform version 3.8 was used for mapping reads to the human hg38 reference and for the base calling of samples FAD24, FAD25 and CTL16 (https://support-docs.illumina.com/SW/DRAGEN_v38/Content/SW/FrontPages/DRAGEN.htm).

Resulting variant calls were annotated using a custom pipeline developed at The Centre for Applied Genomics (TCAG) at SickKids based on ANNOVAR [3]. Structural variants were called using Manta v0.29.6 [4] and LUMPY v0.2.13 [5]. LUMPY calls were genotyped using SVTyper v0.1.2 [6]. Other protocols specific to copy number variants calling from WGS using read depth method were done by the programs ERDS v1.1 and CNVnator v0.3.2 using a window size of 500 bp [7, 8]. Variant prioritization was done on candidate genes for AD (Supplemental Figure X). Population frequency of candidate variants were verified on thousand genomes (1000g phase3;[9] ) and Genome Aggregation Database v2.1.1 (gnomAD; [10]). Mutations and genotypes well known to cause AD in those genes were manually inspected from mapping file of affected and control samples on Integrative Genomic Viewer [11]. Pipeline to identify putative causative variants for Alzheimer's disease (AD) in samples AD1 and AD2 from WGS data: A) Manta [4] and LUMPY [5] algorithms were used to identify structural variants on candidate genes; B) CVNnator [7] and ERDS [12] were also used to identify copy number variants (CNV) as these methods are more accurate, but they do not detect transposable elements (TE), inversions and translocations. C) SpliceAI (https://github.com/Illumina/SpliceAI; [13]) was used to verify if mutations were likely to create an aberrant splicing event. Coding variants on the panel genes identified for samples AD1 and samples AD2 were deemed not likely to be disease causing because population frequency for each was over 30% in genome AD and 1000genomes. Some of those variants were flagged in ClinVar (https://www.ncbi.nlm.nih.gov/clinvar) as benign or likely benign (Table 1). Intronic variants considered rare were not predicted to cause strong splicing effects using SpliceAI (Table 1). The combination of *APOE* SNPs rs7412 and rs429358 resulted in genotype E3/E3 for the two AD samples and control CTL2 and E2/E3 for control CTL1 (Table 2). Both E3/E3 and E2/E3 genotypes are the common genotypes in controls are not associated with increased risk of AD [14]. In AD2 was identified a SNP upstream to *BMI1* (rs17415557, Table 3) previously reported as protective against AD [15]. Other SNPs relevant for AD in *APP*, *PLD3*, *TREM2*, and *TM2D3* were not identified in cell lined derived from affected or control cell individuals (Table 2).

Table 1. Most relevant variants on panel genes identified for AD1, AD2, FAD7, FAD24, and FAD25 samples.

| Sample | VCF[1] | gene | effect | Zygosity[2] | coding position[3] | Protein change prediction | 1000g[4] | gnomAD[4] |
|---|---|---|---|---|---|---|---|---|
| AD2 | 1:227058344:A:G | *PSEN2* | 5'UTR | het | -11265A>G | | 0.540535 | 0.5079 |
| AD1,AD2 | 1:227069677:T:C | *PSEN2* | synonymous | het | 69T>C | (Ala23Ala) | 0.735623 | 0.7699 |
| AD1,AD2 | 1:227069737:C:T | *PSEN2* | synonymous | het | 129C>T | (Asn43Asn) | 0.443291 | 0.4779 |
| AD1,AD2 | 1:227071525:C:T | *PSEN2* | synonymous | het | 261C>T | (His87His) | 0.443291 | 0.4782 |
| AD2 | 6:41250466:T:A | *APOE* | nonsynonymous | het | 42C>G | (Asn14Lys) | 0.626198 | 0.696 |
| AD1 | 11:121403229:T:C | *SORL1* | synonymous | hom | T1653T>C | (Ala551Ala) | 0.985224 | 0.9867 |
| AD1 | 11:121426042:G:A | *SORL1* | splicing region | het | 2571+15G>A | | 0.0439297 | 0.0332 |
| AD1,AD2 | 11:121437819:C:G | *SORL1* | nonsynonymous | hom | 3220C>G | (Gln1074Glu) | 0.984824 | 0.9851 |
| AD1,AD2 | 11:121445084:T:C | *SORL1* | splicing region | hom | 3460+12T>C | | 0.984824 | 0.985 |
| AD1 | 11:121456962:C:T | *SORL1* | synonymous | hom | 3738C>T | (Asn1246Asn) | 0.407348 | 0.5687 |
| AD1,AD2 | 11:121460846:C:T | *SORL1* | synonymous | ref | 4176C>T | (Asn1392Asn) | 0.0840655 | 0.0669 |
| AD1 | 11:121475922:T:A | *SORL1* | synonymous | ref | 4752T>A | (Asn1584Asn) | 0.396166 | 0.3023 |
| AD1 | 11:121491782:G:A | *SORL1* | nonsynonymous | hom | 5899G>A | (Val1967Ile) | 0.979433 | 0.9853 |
| AD2 | 11:121367626:T:C | *SORL1* | synonymous | het | 807T>C | (His269His) | 0.44369 | 0.4923 |

[1] VCF- genomic position on reference hg19. Chromosome: position: wildtype: variant
[2]Zygosity in the first sample – heterozygous (het), homozygous (hom)
[3] *PSEN2* (NM_000447), *SORL1* (NM_003105), *PSEN1* (NM_000021)
[4] Population frequency across all populations of the database.

Table 2. Genotype of known variants associated with late onset Alzheimer's disease in affected and control samples.

| Gene | dbSNP | chr | Position[a] | reference | AD1 | AD2 | CTL1 | CTL 2 | CTL17 | CTL16 |
|---|---|---|---|---|---|---|---|---|---|---|
| *APP* | rs63750847[1] | 21 | 27269932 | [16] | G/G | G/G | G/G | G/G | G/G | G/G |
| *APOE* | rs7412 | 19 | 45412079 | [14] | C/C | C/C | C/T | C/C | C/C | C/C |
| *APOE* | rs429358 | 19 | 45411941 | [14] | T/T | T/T | T/T | T/T | T/T | T/T |
| *PLD3* | rs145999145 | 19 | 40877595 | [17] | G/G | G/G | G/G | G/G | G/G | G/G |
| *TREM2* | rs75932628 | 6 | 41129252 | [16] | C/C | C/C | C/C | C/C | C/C | C/C |
| *TM2D3* | rs139709573 | 15 | 102186966 | [18] | G/G | G/G | G/G | G/G | G/G | G/G |
| *BMI1* | rs17415557 | 10 | 22600627 | [15] | T/T | T/G | T/T | T/T | T/T | T/T |

[a] reference hg19
[1] The presence of A or AA protective against AD.

### Supplementary Method References

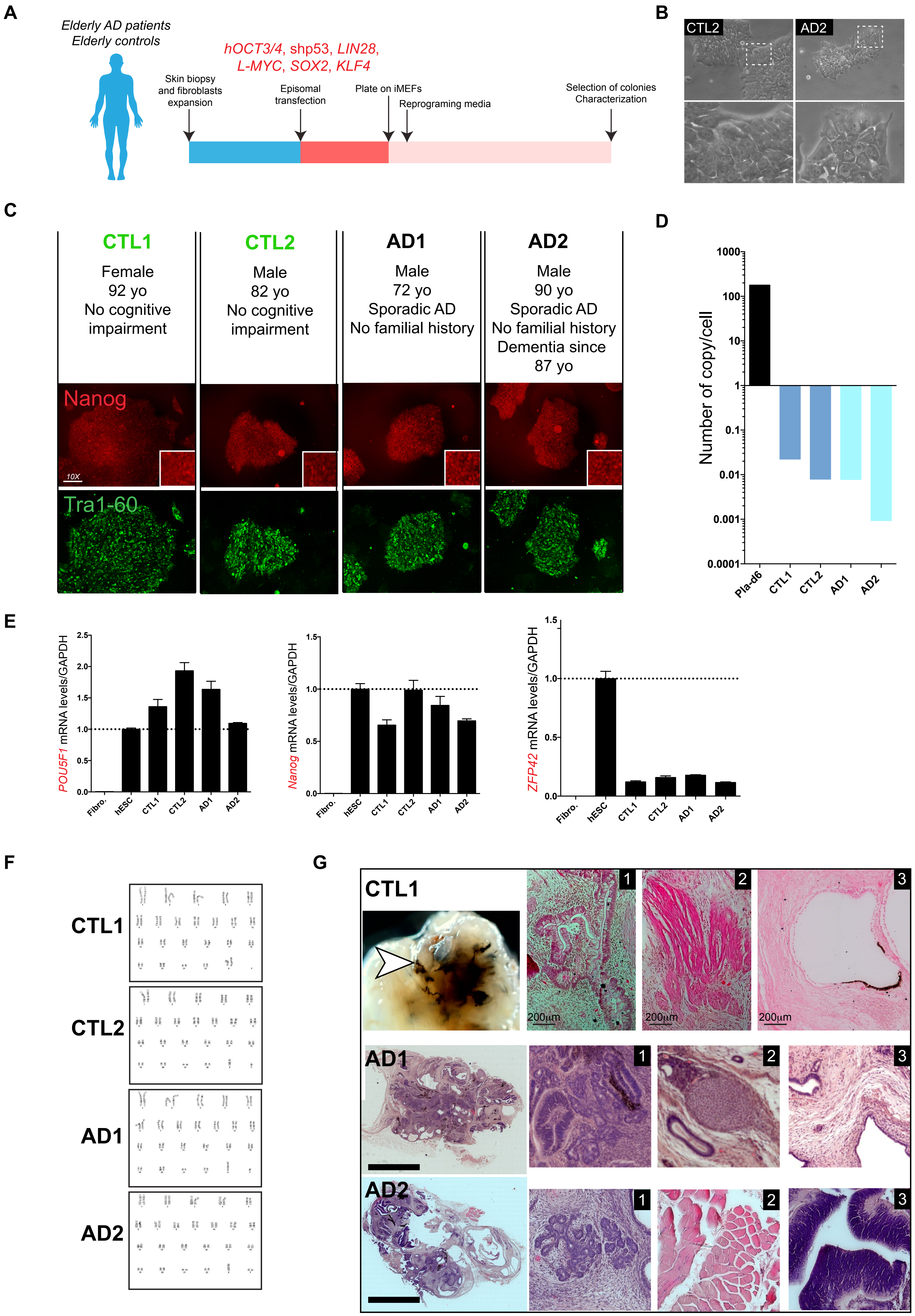

A
Elderly AD patients
Elderly controls
hOCT3/4, shp53, LIN28,
L-MYC, SOX2, KLF4
Skin biopsy
and fibroblasts
expansion
Episomal
transfection
Plate on iMEFs
Reprograming media
Selection of colonies
Characterization
B
CTL2
AD2
C
CTL1
Female
92 yo
No cognitive
impairment
CTL2
Male
82 yo
No cognitive
impairment
AD1
Male
72 yo
Sporadic AD
No familial history
AD2
Male
90 yo
Sporadic AD
No familial history
Dementia since
87 yo
Nanog
10X
Tra1-60
D
Number of copy/cell
1000
100
10
1
0.1
0.01
0.001
0.0001
Pla-d6
CTL1
CTL2
AD1
AD2
E
POU5F1 mRNA levels/GAPDH
Nanog mRNA levels/GAPDH
ZFP42 mRNA levels/GAPDH
Fibro.
hESC
CTL1
CTL2
AD1
AD2
F
CTL1
CTL2
AD1
AD2
G
CTL1
AD1
AD2
200µm
1
2
3


Figure S1

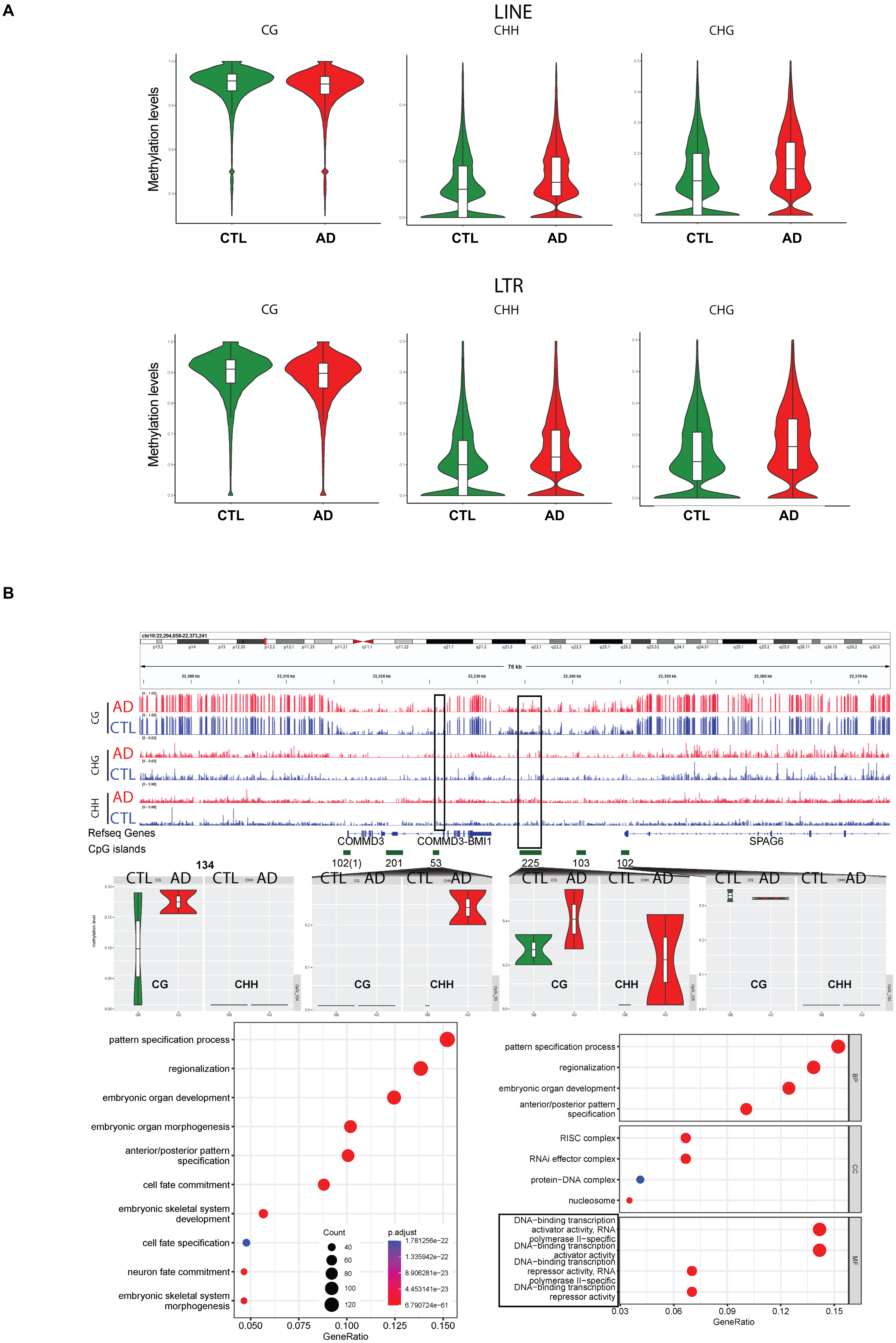

A
LINE
CG
CHH
CHG
Methylation levels
CTL
AD
LTR
B
chr10:22,294,658-22,373,241
78 kb
22,300 kb
22,310 kb
22,320 kb
22,330 kb
22,340 kb
22,350 kb
22,360 kb
22,370 kb
Refseq Genes
CpG islands
COMMD3
COMMD3-BMI1
SPAG6
102(1)
201
53
225
103
102
134
pattern specification process
regionalization
embryonic organ development
embryonic organ morphogenesis
anterior/posterior pattern specification
cell fate commitment
embryonic skeletal system development
cell fate specification
neuron fate commitment
embryonic skeletal system morphogenesis
Count
40
60
80
100
120
p.adjust
1.781256e−22
1.335942e−22
8.906281e−23
4.453141e−23
6.790724e−61
0.050
0.075
0.100
0.125
0.150
GeneRatio
RISC complex
RNAi effector complex
protein−DNA complex
nucleosome
DNA−binding transcription activator activity, RNA polymerase II−specific
DNA−binding transcription activator activity
DNA−binding transcription repressor activity, RNA polymerase II−specific
DNA−binding transcription repressor activity
BP
CC
MF
0.03
0.06
0.09
0.12
0.15


Figure S2

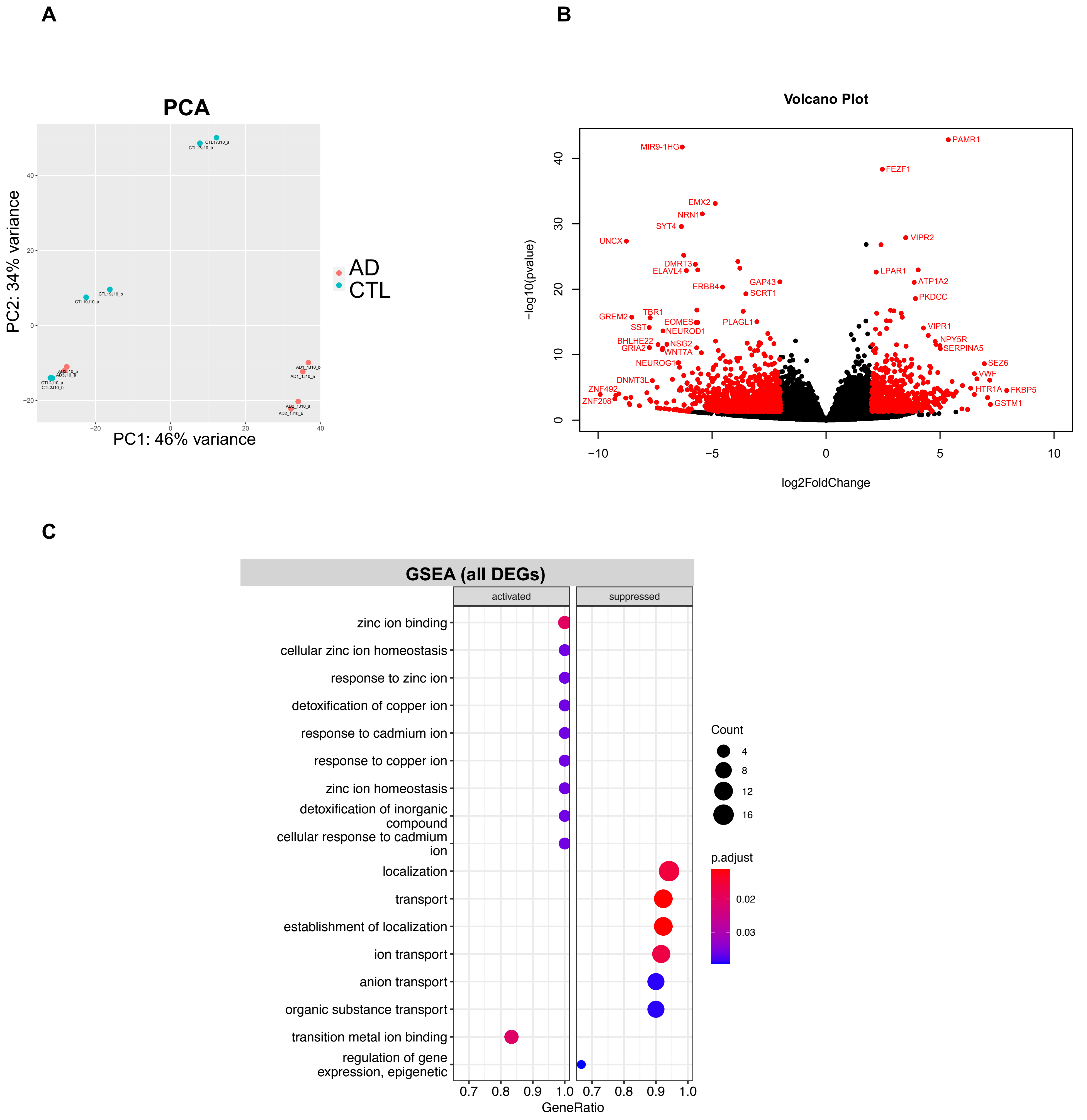
A
PCA
PC2: 34% variance
PC1: 46% variance
AD
CTL
B
Volcano Plot
−log10(pvalue)
log2FoldChange
MIR9-1HG
PAMR1
FEZF1
EMX2
NRN1
SYT4
UNCX
VIPR2
DMRT3
ELAVL4
LPAR1
ATP1A2
GAP43
ERBB4
SCRT1
PKDCC
GREM2
TBR1
EOMES
PLAGL1
SST
NEUROD1
VIPR1
BHLHE22
NSG2
NPY5R
GRIA2
WNT7A
SERPINA5
NEUROG1
SEZ6
VWF
DNMT3L
ZNF492
HTR1A
FKBP5
ZNF208
GSTM1
C
GSEA (all DEGs)
activated
suppressed
zinc ion binding
cellular zinc ion homeostasis
response to zinc ion
detoxification of copper ion
response to cadmium ion
response to copper ion
zinc ion homeostasis
detoxification of inorganic compound
cellular response to cadmium ion
localization
transport
establishment of localization
ion transport
anion transport
organic substance transport
transition metal ion binding
regulation of gene expression, epigenetic
GeneRatio
Count
p.adjust

Figure S3

A

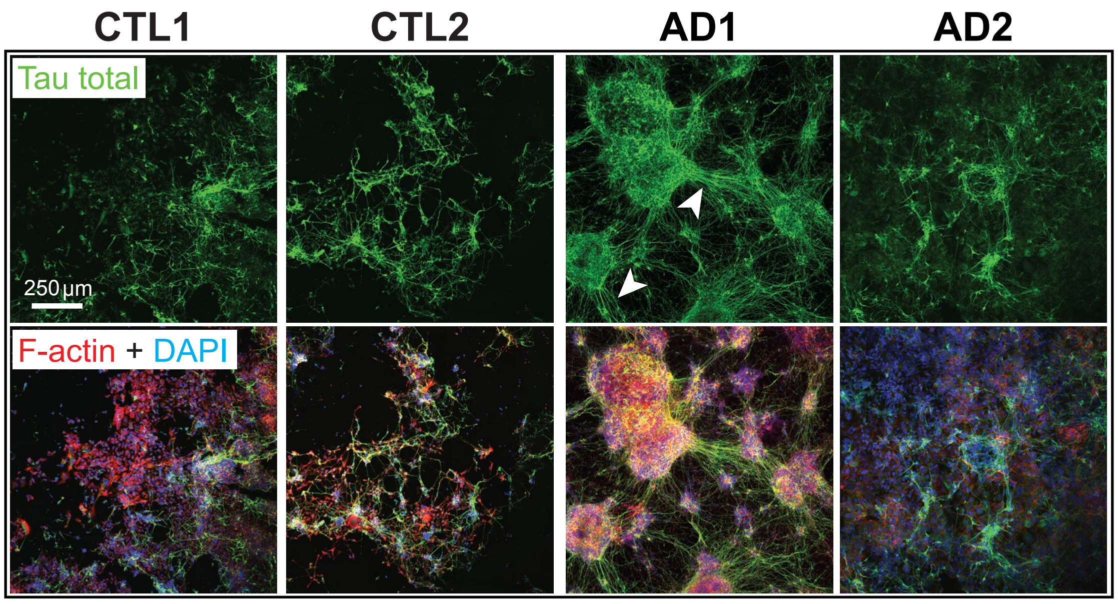
CTL1
CTL2
AD1
AD2
Tau total
250 µm
F-actin + DAPI

B

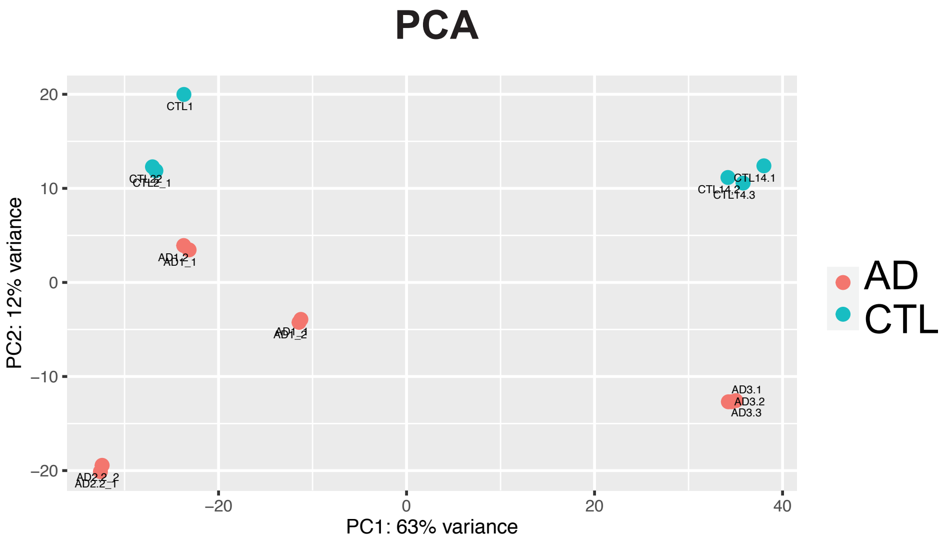
PCA
PC2: 12% variance
PC1: 63% variance
AD
CTL
CTL1
AD3.1
AD3.2
AD3.3
CTL14.1
CTL14.3

Figure S4

**A**

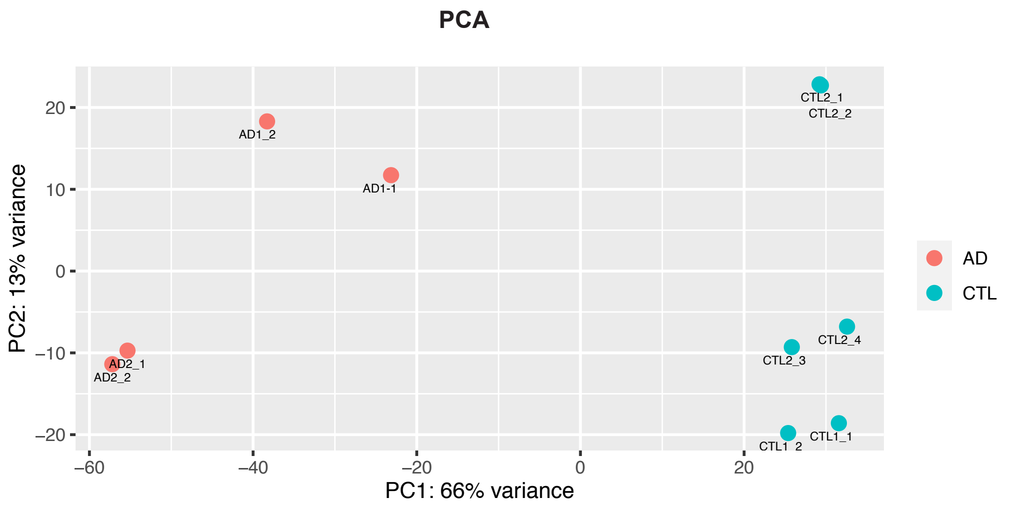


**B**

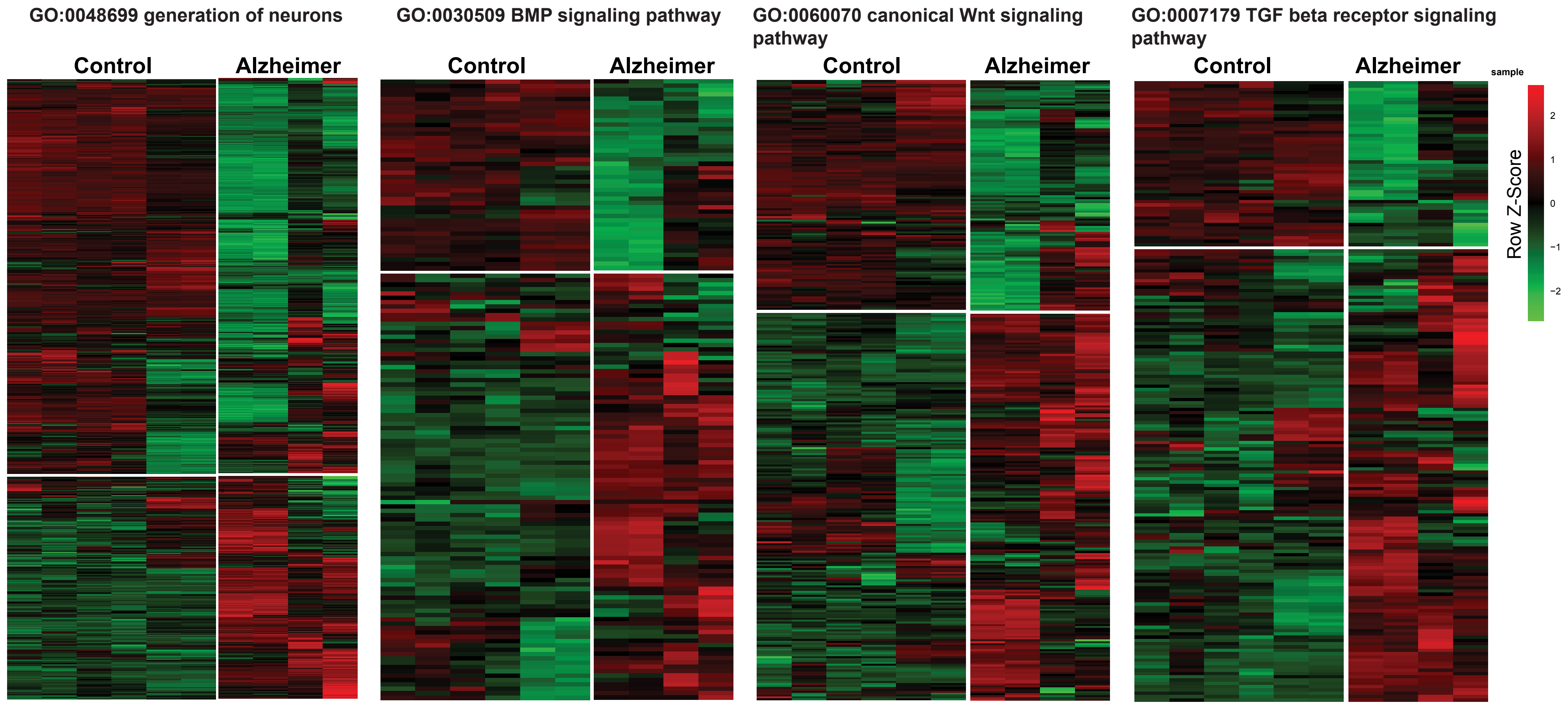


**C**

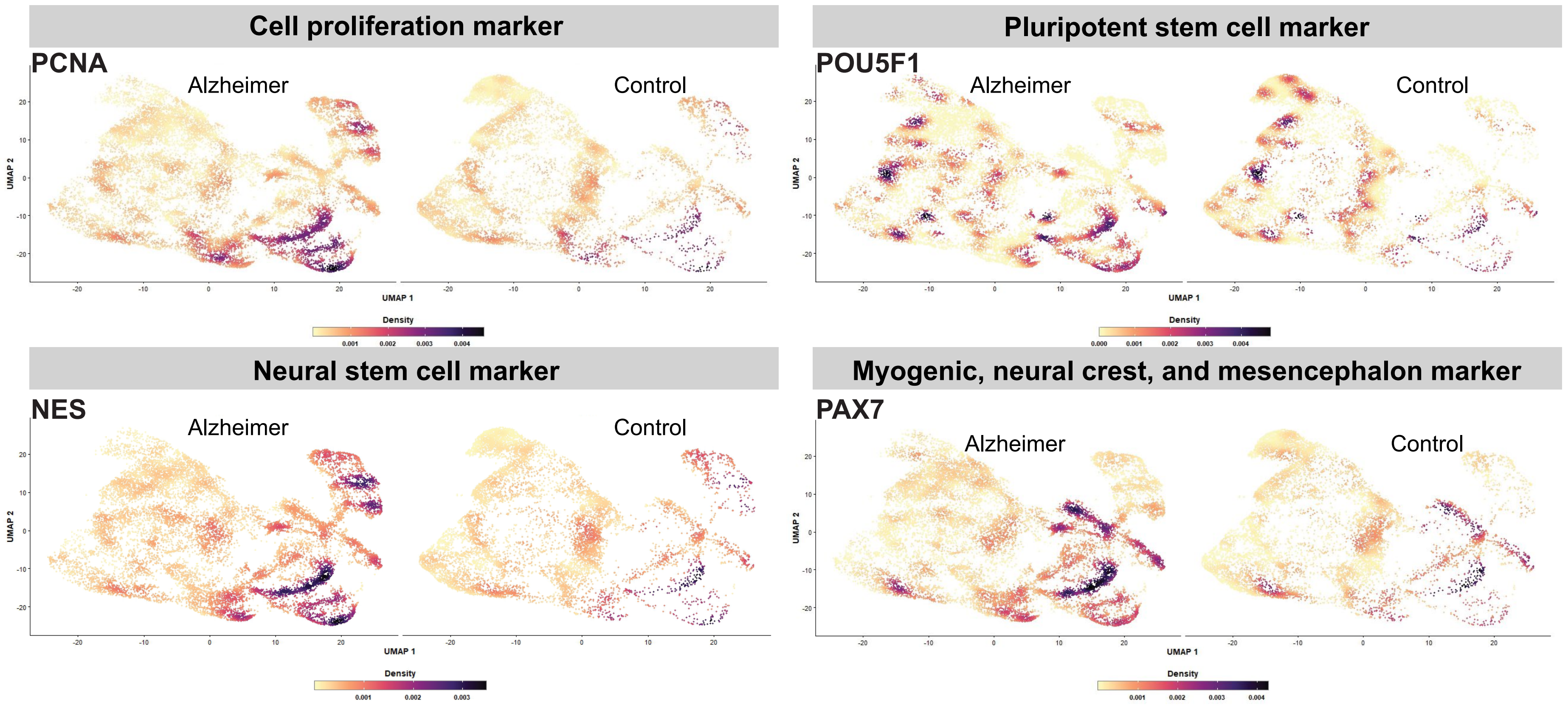


Figure S5

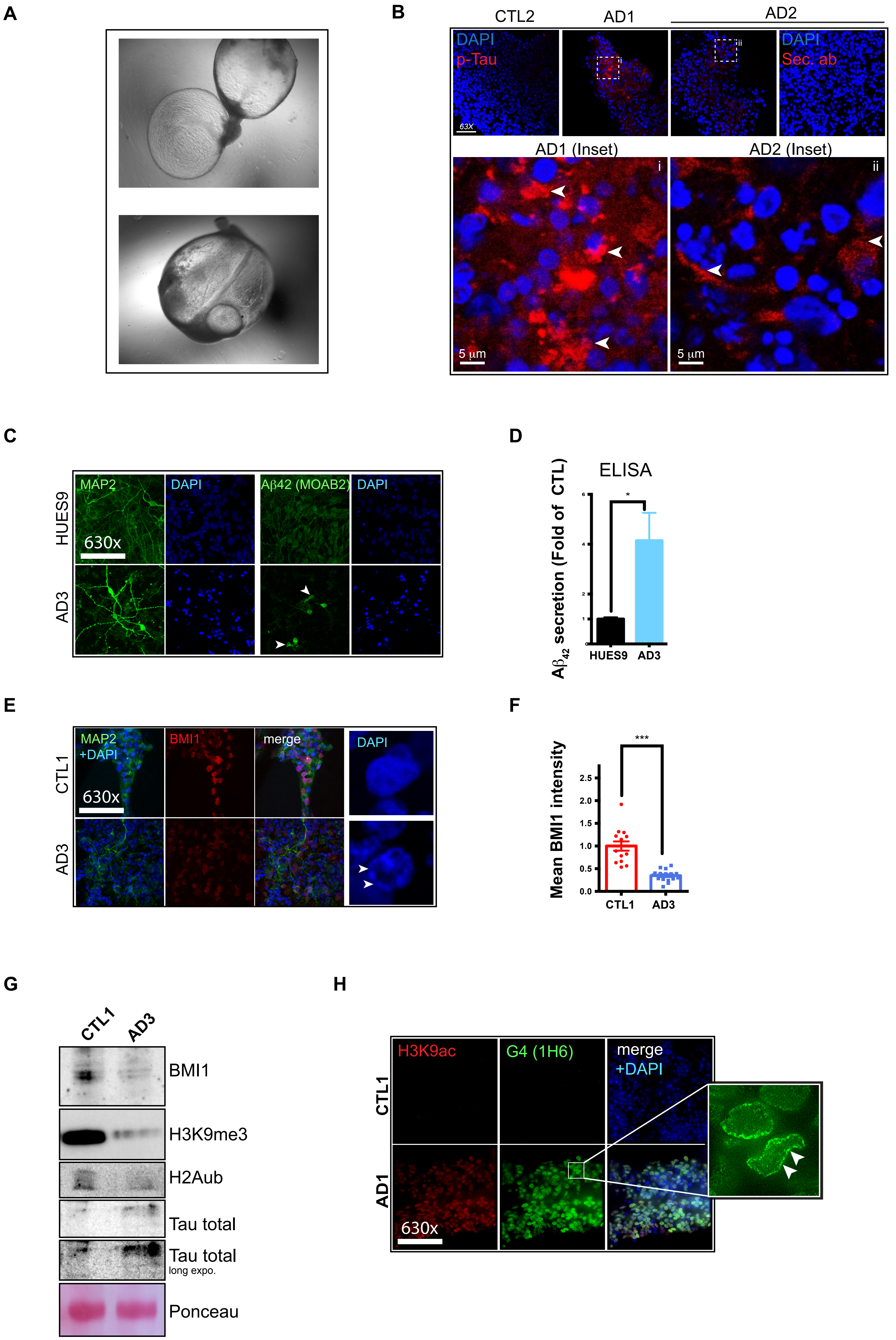
A
B
CTL2
AD1
AD2
DAPI
p-Tau
DAPI
Sec. ab
63X
AD1 (Inset)
AD2 (Inset)
i
ii
5 µm
5 µm
C
MAP2
DAPI
Aβ42 (MOAB2)
DAPI
HUES9
AD3
630x
D
ELISA
Aβ42 secretion (Fold of CTL)
HUES9
AD3
E
MAP2
+DAPI
BMI1
merge
DAPI
CTL1
AD3
630x
F
Mean BMI1 intensity
***
2.5
2.0
1.5
1.0
0.5
0.0
CTL1
AD3
G
CTL1
AD3
BMI1
H3K9me3
H2Aub
Tau total
Tau total
long expo.
Ponceau
H
H3K9ac
G4 (1H6)
merge
+DAPI
CTL1
AD1
630x

Figure S6

### SUPLEMENTARY FIGURE LEGENDS

**Figure S1. Generation of iPSCs from control and AD patient's fibroblasts**

(A) Schematic overview for the reprogramming of dermal fibroblasts into induced pluripotent stem cells (iPSCs).

(B) Phase contrast representative images of one control iPSC line (CTL2) and one sporadic AD iPSC line (AD2) adapted to Matrigel culture conditions. The insets exemplify the high nuclear/cytoplasm ratio, characteristic of pluripotency.

(C) Immunofluorescence analysis on CTL1, CTL2, AD1 and AD2 iPSC line colonies for pluripotency markers Nanog and Tra1-60. Note the nuclear localization of Nanog in the insets. Scale bar: 40µm.

(D) Quantification of episomal integrations in CTL1, CTL2, AD1 and AD2 iPSC lines by quantitative genomic PCR. Fibroblasts after 6 days of reprogramming were used as positive control (Pla-d6).

(E) *POU5F1* (OCT3/4), *NANOG* and *ZFP42* (REX1) gene expression levels in qPCR normalized to GAPDH on CTL1, CTL2, AD1 and AD2 iPSC lines. Dermal fibroblasts (Fibro.) and human embryonic stem cell line H9 (hESC) were used as negative and positive controls respectively ($n = 3$ for each line). All values are mean ± SEM.

(F) Karyotyping analyses on CTL1, CTL2, AD1 and AD2 iPSC lines by G-banding.

(G) Teratoma formation after injection of CTL1, AD1 and AD2 iPSC lines under the skin of NOD/SCID mice. Hematoxylin and Eosin histology shows the formation of glandular epithelium (endoderm) (1), skeletal muscle or cartilage (mesoderm) (2), and retinal pigment epithelium or neuroepithelium (ectoderm) (3).

**Figure S2. Analysis by bWGS of CTL and AD iPSC lines**

(A) Violon plot of CG, CHH and CHG DNAme between CTL1/CTL2 and AD1/AD2 iPSC lines at LINE and LTR elements. Note the high CG DNAme in both CTL and AD iPSC lines at LINEs and LTRs.

(B) Physical map of the *COMMD3* and *COMMD3-BMI1* locus, with represented CG, CHG, and CHH DMRs between CTLs and ADs. Middle panel: No DMRs were observed for the entire locus between CTLs and ADs. CG and CHH hypermethylation was however observed in AD iPSCs at CpG Islands 134, 53, and 225 (in red). Bottom panel: Gene ontology analysis of CpG Island-associated genes with CG and CHH hypermethylation in AD iPSCs.

**Figure S3. Differentiation of iPSCs into neuroepithelial cells**

(A) Principal component analysis (PCA) of CTL and AD NE cells at DIV10.

(B) Volcano plot of all genes expressed in CTL and AD NE cells. Significant DEGs at more than 2 folds (red dots).

(C) GSEA of DEGs between CTL and AD NE cells, showing activated or suppressed pathways in AD.

**Figure S4. Differentiation of iPSCs into immature cortical neurons**

(A) Confocal immunofluorescence analysis images. Neuronal cultures were labeled at DIV30 with antibodies against Tau (green) and F-actin (Phalloidin-red). Note the accumulation of Tau and F-actin in the AD1 culture (arrowheads). Scale bar: 250μm.

(B) Principal component analysis (PCA) of CTL and AD immature neurons at DIV30.

**Figure S5. Differentiation of iPSCs into mature cortical neurons**

(A) Principal component analysis (PCA) of CTL and AD neuronal cultures at DIV60. Note segregation of the CTL and AD groups.

(B) Heatmap of all DEGs between CTL and AD iPSC lines for 4 distinct biological pathways, with a cutoff of at least 2 folds. Note suppression of neurogenesis and activation of WNT, BMP and TGFb signaling pathways in AD cultures.

(C) UMAP plot comparison between CTL and AD neuronal cultures at DIV60 for 4 different markers. Note the cell proliferation, stem cell, and non-neuronal gene expression phenotype of AD cultures.

**Figure S6. Alzheimer-like pathology in AD brain organoids and neuronal cultures**

(A) Representative bright field images of floating brain organoids after 4 months in culture.

(B) Immunofluorescence on sections of CTL and AD brain organoids. p-Tau accumulates in AD1 and AD2 brain organoids but not in CTL2 brain organoids at 4 months of differentiation. Sections from the AD2 brain organoids were also labeled only with the secondary antibody to test for possible non-specific fluorescence (Sec. ab). Scale bar: 40μm. The white arrowheads in the high magnification images (inset) indicate p-Tau deposits resembling Tau tangles. Scale bar: 5μm.

(C-F) Characterization of the AD3 iPSC line after neuronal differentiation for 60 days. Note the abnormal accumulation and secretion of Aβ42 in AD3 cultures when compared to CTL as revealed by immunofluorescence (C) and ELISA (D). BMI1 protein levels were also reduced by 50-60% in AD3 neurons, as showed by quantitative immunofluorescence (E-F). All values are mean ± SEM. (***) P< 0.001 by Student's unpaired t-test. Scale bar: 40μm.

(G) Western blot analysis of CTL1 and AD3 neuronal cultures at DIV60, showing reduced BMI1, H3K9me3 and H2Aub levels in AD3, but with increased Tau levels. Ponceau red was used for normalization of total protein concentration.

(H) Immunofluorescence analysis of CTL1 and AD1 neuronal cultures at DIV60 showing increased H3K9ac and G4s (with the 1H6 antibody) levels in AD1. Note the peri-nuclear accumulation of G4s in AD1 cells (arrowheads). Scale bar: 40μm.
